## Supplement file for "Interpretable Online Network Dictionary Learning for Inferring Long-Range Chromatin Interactions"

#### 1 Motivation

Dictionary learning (DL), a form of nonnegative matrix factorization (MF), has been widely used in the analysis of biological data. However, *efficient*, and *biologically interpretable* computational methods for analyzing long-distance multiplexed chromatin interactions at a single-cell level are still lacking. This gap exists primarily because classical DL methods are not tailored for network data analysis. Furthermore, these interactions cannot be easily visualized or predicted via classical clustering approaches. This issue is best illustrated by Figure 1, where a part of the contact map contains three hidden clusters, colored red, green, and blue [1]. When using a linear chromatin order, the particular structure of the clusters is not observable. By rearranging the rows/columns, the cluster structure becomes apparent within the adjacency matrix. To mit-

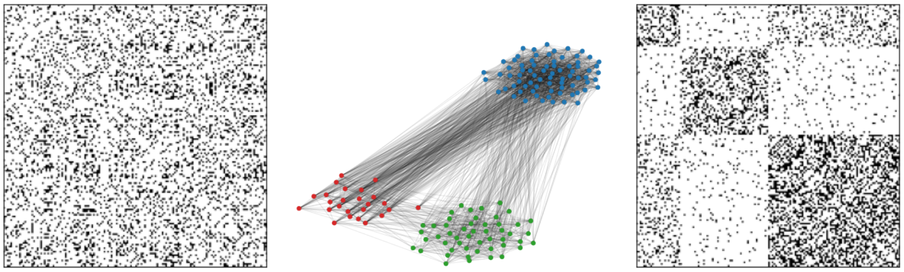

(a) (b) (c)

Figure 1: **(a)** Observed adjacency matrix of a three-cluster model, where points are arranged in linear order with dense interactions existing both at short- and long-range. **(b)** The underlying cluster structure. **(c)** The reordered adjacency matrix that reveals all interaction classes.

igate this issue, we propose a novel online convex network dictionary learning algorithm (online cvxNDL) that imposes "convexity" constraints on the sampled subgraph patterns to address the issue of interpretability. Furthermore, due to its online nature, it scales to large graph-structured datasets. The detailed algorithmic implementations are described in the next section.

### 2 Algorithmic Details

The algorithms presented in this section describe the detailed steps of implementation outlined in the Methods Section.

#### 2.1 MCMC Sampling of Subnetworks

We use the MCMC sampling in conjunction with subnetwork sampling to generate online samples. We seek samples in the form of subnetworks induced by $k$ nodes in the original input network $\mathcal{G}$ such that these subnetworks contain the template $F$ topology. Given an input network $\mathcal{G} = (V, \mathbf{A})$ and a template network $F = ([k], \mathbf{A}_F)$, we define a set of homomorphisms as a vector of the form (with the assumption that $0^0 = 1$):

$$\mathtt{Hom}(F, \mathcal{G}) = \left\{ \underline{x} : [k] \to [n] \,\middle|\, \prod_{1 \le i,j \le k} \mathbf{A}[\underline{x}[i], \underline{x}[j]]^{\mathbf{A}_F[i,j]} = 1 \right\}.$$

Algorithm 1 outlines how to use rejection sampling to obtain one homomorphism $\underline{x}$ (an illustrative example is presented in Figure 1(d) in the main text). In this work, we use a $k$-path as the template network, where a $k$-path represents a directed path from node 1 to $k$. Paths serve as a simple and natural choice for networks containing inherent long paths, such as chromatin interaction networks, where most contact measurements are due to proximity in the linear chromosome order.

**Algorithm 1** Rejection Sampling of Homomorphisms

**input:** Network $\mathcal{G} = ([n], \mathbf{A})$, template $F = ([k], \mathbf{A}_F)$ (under the assumption that there exists at least one homomorphism $F \to \mathcal{G}$).
**while** true **do**
  Sample $\underline{x} = (\underline{x}[1], \underline{x}[2], \ldots, \underline{x}[k]) \in [n]^k$ so that $\underline{x}[i]$'s are i.i.d.
  **if** $\prod_{1 \le i,j \le k} \mathbf{A}[\underline{x}[i], \underline{x}[j]]^{\mathbf{A}_F[i,j]} > 0$ **then**
    **break**
  **end if**
**end while**
**return** A homomorphism $\underline{x} : F \to \mathcal{G}$.

While we can find different homomorphisms from the input $\mathcal{G}$ by iteratively executing Algorithm 1, this method is computationally expensive. To efficiently generate a sequence of sample adjacency matrices $\mathbf{A}_{\underline{x}_t}$ from $\mathcal{G}$, the MCMC sampling algorithm gradually changes the sampled subnetwork based on previous samples as described in Algorithm 2. An illustrative example is shown in Figure 1(e) in the main text. This sampling algorithm was introduced in [2, 3].

**Algorithm 2** The MCMC Sampling Algorithm

**input:** Network $\mathcal{G} = ([n], \mathbf{A})$, template $F = ([k], \mathbf{A}_F)$, and one homomorphism $\underline{x} : F \to \mathcal{G}$.
Sample $v \in \text{Neighbor}(\underline{x}[1])$ with probability $P(v) = \frac{1}{\mathcal{N}[\underline{x}[1]]}$.
Compute the acceptance probability
$\beta = \min\left\{ \frac{\sum_{c\in[n]} A^{k-1}[v,c]}{\sum_{c\in[n]} A^{k-1}[\underline{x}[1],c]} \frac{\sum_{c\in[n]} A[\underline{x}[1],c]}{\sum_{c\in[n]} A[v,c]}, 1 \right\}$.
Sample $u$ uniformly at random from $[0, 1]$.
**if** $u < \beta$ **then**
    $\underline{x}'[1] = v$
**else**
    $\underline{x}'[1] = \underline{x}[1]$
**end if**
**for** $s = 2, 3, \ldots, k$ **do**
    Sample $w \in [n]$ with probability $P_s(w) = \frac{\mathbf{A}[\underline{x}'[s-1],w]}{\sum_{c\in V} \mathbf{A}[\underline{x}'[s-1],c]}$.
    $\underline{x}'[s] = w$
**end for**
**return** New homomorphism $\underline{x}' : F \to \mathcal{G}$.

#### 2.2 Online Convex NDL (online cvxNDL)

Our online cvxNDL algorithm consists of two parts: initialization and iterative optimization. For initialization, we compute an initial choice for the dictionary elements $\mathbf{D}_0$ and initialize the representative regions $\hat{\mathbf{X}}_0^{(j)}$, $\forall j \in [K]$ using i.i.d. sampling of homomorphisms (Algorithm 3). Note that we use i.i.d. sampling of homomorphisms only during the initialization step, and MCMC sampling afterwards. Upon initialization, we iteratively optimize the dictionary and the representative regions in the next phase (Algorithm 4). The output of the latter algorithm is the final dictionary $\mathbf{D}_T$ and the corresponding representative regions for all dictionary elements $\hat{\mathbf{X}}_T^{(j)}$, $\forall j \in [K]$. Due to the added convexity constraint, each dictionary element $\mathbf{D}_T[:, j]$ at the final step $T$ has the following interpretable form:

$$\mathbf{D}_T[:, j] = \sum_{i\in[N_j]} w_{j,i} \hat{\mathbf{X}}_T^{(j)}[:, i], \text{ s.t. } \sum_{i\in[N_j]} w_{j,i} = 1, w_{j,i} \geq 0, i \in [N_j], j \in [K].$$

The weight $w_{j,i}, i \in [N_j]$ is the convex coefficient of the $i^{\text{th}}$ representative of dictionary element $\mathbf{D}_T[:, j]$.

**Algorithm 3** Initialization

1: **input:** Use rejection sampling in Algorithm 1 to sample i.i.d homomorphisms $\underline{x}_1, \underline{x}_2, \ldots, \underline{x}_N$.

2: For each homomorphism, define an adjacency matrix such that: $\mathbf{A}_{\underline{x}_i}[a, b] = \mathbf{A}[\underline{x}_i[a], \underline{x}_i[b]]$. Flatten the adjacency matrices into vectors: $\underline{x}_1, \underline{x}_2, \ldots, \underline{x}_N$, $\underline{x}_i \in \mathbb{R}^d, d = k^2$ and collect them in $\hat{\mathbf{X}} \in \mathbb{R}^{d \times N}$.

3: Run $K$-means on $\hat{\mathbf{X}}$ to generate the cluster indicator matrix $\mathbf{H} \in \{0, 1\}^{N \times K}$ and determine the initial cluster sizes (subsequent representative set sizes) $N_i, i \in [K]$.

4: Compute $\mathbf{D}_0$ and $\hat{\mathbf{X}}_0^{(i)} \in \mathbb{R}^{d \times N_i}, \ \forall i \in [K]$, according to:

$$\mathbf{D}_0 = \hat{\mathbf{X}} \ \mathbf{H} \ \mathrm{diag}(1/N_1, \ldots, 1/N_K)$$

and summarize the initial representative sets of the clusters into matrices $\hat{\mathbf{X}}_0^{(i)}$, $i = [K]$.

5: **return** $\mathbf{D}_0$, $\{\hat{\mathbf{X}}_0^{(i)}\}_{i \in [K]}$.

**Algorithm 4** Online cvxNDL

**input:** Network $\mathcal{G} = ([n], \mathbf{A})$, template $F = ([k], \mathbf{A}_F)$, a parameter $\lambda \in \mathbb{R}$, max number of iterations $T$, and number of dictionary elements $K$.

**initialization:** Compute $\mathbf{D}_0$, $\{\hat{\mathbf{X}}_0^{(i)}\}_{i \in [K]}$ using Algorithm 3. Set $\mathbf{A}_0 = \mathbf{0}$, $\mathbf{B}_0 = \mathbf{0}$.

**for** $t = 1$ to $T$ **do**

MCMC sample a homomorphism $\underline{x}_t$ (Algorithm 2). Find its adjacency matrix $\mathbf{A}_{\underline{x}_t}[a, b] = \mathbf{A}[\underline{x}_t[a], \underline{x}_t[b]]$ and flatten it to $\underline{x}_t$.

Update $\mathbf{\Lambda}_t$ according to:

$$\mathbf{\Lambda_t} = \arg\min_{\mathbf{\Lambda} \in \mathbb{R}^{K \times 1}} \frac{1}{2} \left\| \underline{x}_t - \mathbf{D}_{t-1} \mathbf{\Lambda} \right\|_2^2 + \lambda \left\| \mathbf{\Lambda} \right\|_1 . \tag{1}$$

Set $\mathbf{A}_t = \frac{1}{t}((t-1)\mathbf{A}_{t-1} + \mathbf{\Lambda}_t \mathbf{\Lambda}_t^T)$ and $\mathbf{B}_t = \frac{1}{t}((t-1)\mathbf{B}_{t-1} + \underline{x}_t \; \mathbf{\Lambda}_t^T)$.

Choose the index of the basis $i_t$ to be updated according to $i_t = \arg\max_{j \in [k]} \mathbf{\Lambda}_t[j]$

Generate the augmented representative regions $\{\hat{\mathbf{Y}}_t^l\}_{l \in [N_{i_t}] \cup \{0\}}$:

$$\begin{aligned} \hat{\mathbf{Y}}_t^0 &= \hat{\mathbf{X}}_{t-1}^{i_t} \\ \{\hat{\mathbf{Y}}_t^l\}_{l \in [N_{i_t}]} : \; \hat{\mathbf{Y}}_t^l[j] &= \begin{cases} \hat{\mathbf{X}}_{t-1}^{i_t}[j], & \text{if } j \in [N_i] \setminus l \\ \underline{x}_t, & \text{if } j = l. \end{cases} \end{aligned} \tag{2}$$

Update $\{\hat{\mathbf{X}}_i^t\}_{i \in [K]}$ and $\mathbf{D}_t$ by executing the following two steps

- Compute $l^\star, \hat{\mathbf{D}}^\star$ by solving the optimization problems:

$$l^\star, \hat{\mathbf{D}}^\star = \arg\min_{\substack{l, \; \mathbf{D} \;\; \text{s.t.} \\ \mathbf{D}[j] \in \text{cvx}\{\hat{\mathbf{X}}_{t-1}^j\} \;\, j \neq i_t, \\ \mathbf{D}[i_t] \in \text{cvx}\{\hat{\mathbf{Y}}_t^l\}}} \frac{1}{2} \operatorname{Tr}(\mathbf{D}^T \mathbf{D} \mathbf{A}_t) - \operatorname{Tr}(\mathbf{D}^T \mathbf{B}_t).$$

- Set

$$\begin{aligned} \hat{\mathbf{X}}_t^i &= \begin{cases} \hat{\mathbf{Y}}_t^{l^\star}, & \text{if } i = i_t \\ \hat{\mathbf{X}}_{t-1}^i, & \text{if } i \in [K] \setminus i_t, \end{cases} \\ \mathbf{D}_t &= \hat{\mathbf{D}}^\star. \end{aligned}$$

**end for**

**return** $\mathbf{D}_T$, $\hat{\mathbf{X}}_T^{(i)}$, $\forall i \in [K]$.

### 3 Synthetic Data Analysis

We tested our online cvxNDL method on a network (graph) generated by Stochastic Block Model (SBM) [1], containing 150 nodes with 3 clusters of size $25, 50, 75$. Due to the small size of the synthetic set, we fixed the number of dictionary elements to $K = 6$ and used a path of length 11 as our template. In the initialization step, we sampled 30 subgraphs from the input synthetic data network, with each dictionary element represented by at least 3 representatives. The maximum number of iterations of the online method was set to $1,000$.

We compared online cvxNDL with various baseline methods, including NMF, CMF, and online NDL. The learned dictionary elements for different methods are shown in Figure 2. The dictionary elements in online NDL and online cvxNDL are ordered by their importance score defined as $\gamma(i) = \frac{\mathbf{A}_t[i,i]^2}{\sum_{j\in[K]} \mathbf{A}_t[j,j]^2}$. Each square block in the subplots indicates one dictionary element in the form of an adjacency matrix. The color-shade reflects the values in the adjacency matrix, with black corresponding to 1 (the largest value) and white corresponding to 0 (the smallest value).

From the results, we can see that dictionaries generated using NMF only contain partial interaction structures and are hard to interpret. The two convex methods, CMF and online cvxNDL, contain the template structure in all learned dictionary elements and show stronger off-diagonal connectivity, which is expected as the input data has slightly stronger connections between the first and last cluster than other pairs (See Figure 1). Online NDL dictionary elements represent “a middle ground” between NMF and online cvxNDL. Dictionary elements 2, 0, and 4 resemble those generated by NMF, while dictionary elements 1, 5, and 3 are similar to the ones generated by online cvxNDL, although with weaker connectivity. Also, the importance score distributions of online NDL and online cvxNDL differ substantially. In online NDL, dictionary element 1 in Figure 2 is the dominant component in representations, whereas, in online cxvNDL, the top two dictionary elements (dictionary elements 2 and 5 in 2) share similar scores and the dictionary elements, in general, have a more balanced distribution of importance scores. From the original adjacency, we can see that there are indeed two different connectivity patterns in the network captured by online cvxNDL.

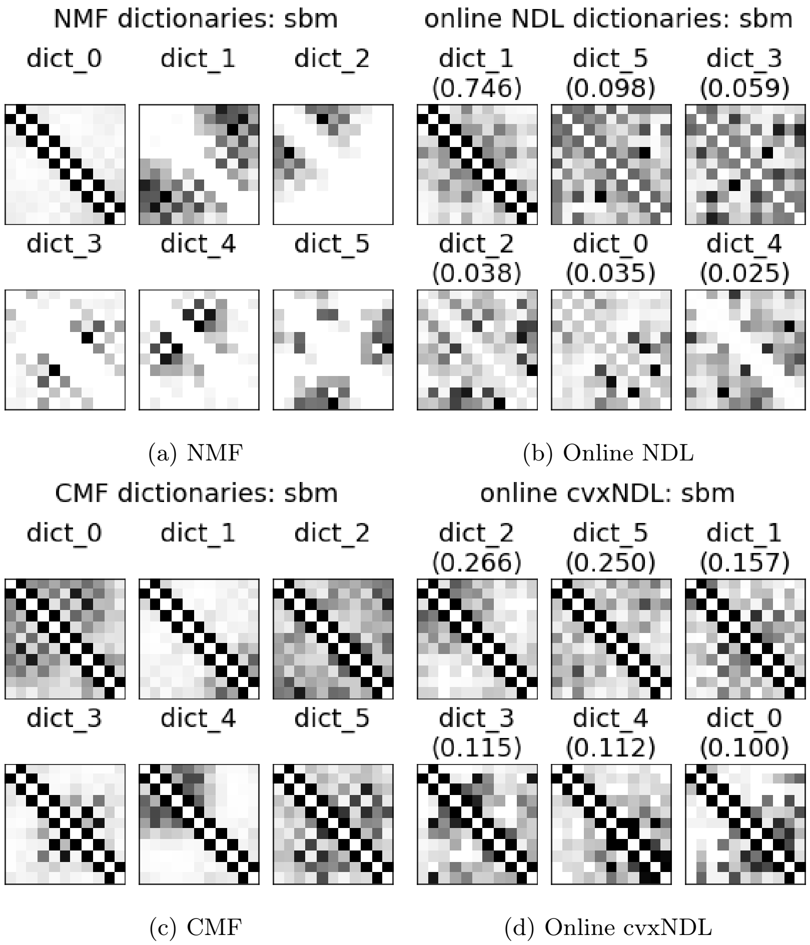


(a) NMF (b) Online NDL

(c) CMF (d) Online cvxNDL

Figure 2: Dictionary elements generated by different methods on an SBM synthetic dataset. Numbers in parenthesis are the importance scores for online NDL and online cvxNDL.

**Reconstruction accuracy:** To validate the reliability of our learned dictionaries for representing the global interactions, we reconstructed the whole graph by aggregating the regenerated subgraphs: $\hat{\mathbf{x}}_i = \mathbf{D}_T \underline{\alpha}_i$ from the same MCMC sampling stream. For each method we selected the top-$m$ edges after aggregation to reconstruct the original adjacency matrix, where $m$ is the number of edges in the original adjacency matrix. The original and the reconstructed adjacency matrices are shown in Figure 7 in the main text. For comparison, we also added the reconstructed adjacency achieved when using random dictionary elements. From the results, we can see that all baseline methods, as well as online cvxNDL, almost perfectly reconstruct the original network, while, clearly random dictionaries do not capture any meaningful information. We also report the average precision recall score for each method, both for synthetic and real datasets as listed in Table 1 in the main text.

### 4 ChIA-Drop Dataset

The preprocessed and binned RNAPII ChIA-Drop data includes $45,938$, $42,292$, $49,072$, and $55,795$ nodes and $36,140$, $28,387$, $53,006$, $45,530$ hyperedges for chromosome chr2L, chr2R, chr3L and chr3R respectively. The size distribution of hyperedges is given in Table 1. The clique-expanded input network has $113,606$, $85,316$, $161,590$, and $143,370$ edges respectively.

Table 1: Number of hyperedges of various sizes observed in the ChIA-Drop data for various chromosomes.

| hyperedge sizes | chr2L | chr2R | chr3L | chr3R |
|---|---|---|---|---|
| 2 | 28373 | 22951 | 42175 | 35585 |
| 3 | 5723 | 4018 | 8103 | 7379 |
| 4 | 1307 | 936 | 1804 | 1700 |
| 5 | 424 | 275 | 533 | 479 |
| 6 | 136 | 94 | 196 | 187 |
| 7 | 60 | 41 | 82 | 69 |
| 8 | 48 | 29 | 38 | 31 |
| 9 | 21 | 15 | 28 | 22 |
| 10 | 8 | 5 | 16 | 7 |
| 11 | 7 | 6 | 9 | 8 |
| 12 | 11 | 2 | 7 | 9 |
| 13 | 5 | 2 | 5 | 7 |
| 14 | 7 | 2 | 2 | 5 |
| 15 | 4 | 2 | 1 | 4 |
| 16 | 3 | 2 | 1 | 4 |
| 17 | 1 | 2 | 2 | 0 |
| 18 | 2 | 1 | 1 | 1 |
| 19 | 0 | 1 | 0 | 0 |
| ≥20 | 1 | 4 | 4 | 7 |

The dictionary elements for each of the 4 chromosomes are presented in Figure 5 in the main text. The density or complexity of dictionary elements, defined as $\rho = \frac{1}{k^2}\sum_{i,j=1}^{k} \mathbf{D}_T[i,j]$, is reported in Table 2 while the median distance of pairwise interacting nodes in all representatives of a dictionary element is reported in Table 3.

Table 2: Density of dictionary elements, reported for all chromosomes.

| Dictionary element | chr2L | chr2R | chr3L | chr3R |
|---|---|---|---|---|
| 1 | 0.146 | 0.158 | 0.168 | 0.161 |
| 2 | 0.188 | 0.165 | 0.156 | 0.157 |
| 3 | 0.134 | 0.185 | 0.141 | 0.140 |
| 4 | 0.220 | 0.147 | 0.159 | 0.179 |
| 5 | 0.145 | 0.146 | 0.142 | 0.139 |
| 6 | 0.132 | 0.297 | 0.148 | 0.173 |
| 7 | 0.162 | 0.189 | 0.191 | 0.184 |
| 8 | 0.158 | 0.184 | 0.164 | 0.147 |
| 9 | 0.148 | 0.136 | 0.210 | 0.183 |
| 10 | 0.177 | 0.166 | 0.168 | 0.157 |
| 11 | 0.220 | 0.261 | 0.163 | 0.161 |
| 12 | 0.168 | 0.162 | 0.145 | 0.157 |
| 13 | 0.204 | 0.203 | 0.186 | 0.142 |
| 14 | 0.225 | 0.142 | 0.148 | 0.205 |
| 15 | 0.142 | 0.229 | 0.262 | 0.163 |
| 16 | 0.173 | 0.184 | 0.143 | 0.205 |
| 17 | 0.189 | 0.263 | 0.127 | 0.224 |
| 18 | 0.161 | 0.219 | 0.152 | 0.251 |
| 19 | 0.182 | 0.159 | 0.183 | 0.242 |
| 20 | 0.187 | 0.156 | 0.170 | 0.193 |
| 21 | 0.231 | 0.157 | 0.199 | 0.126 |
| 22 | 0.143 | 0.195 | 0.165 | 0.150 |
| 23 | 0.162 | 0.201 | 0.134 | 0.175 |
| 24 | 0.223 | 0.141 | 0.167 | 0.212 |
| 25 | 0.167 | 0.212 | 0.140 | 0.208 |

Table 3: Median distance of pairwise interacting nodes within each dictionary element and for each chromosome.

| dictionary element | chr2L | chr2R | chr3L | chr3R |
|---|---|---|---|---|
| 1 | 10758 | 6738 | 7328 | 14753 |
| 2 | 8523 | 7688 | 12934 | 14760 |
| 3 | 9906 | 8759 | 9539 | 12666 |
| 4 | 8354 | 7158 | 12690 | 11748 |
| 5 | 9847 | 7651 | 10412 | 13674 |
| 6 | 8547 | 6953 | 10608 | 15598 |
| 7 | 10024 | 9383 | 11994 | 13498 |
| 8 | 8870 | 9226 | 10399 | 12830 |
| 9 | 10692 | 7085 | 14414 | 12493 |
| 10 | 11220 | 6414 | 9466 | 11930 |
| 11 | 10455 | 10711 | 10130 | 11421 |
| 12 | 8488 | 7656 | 11694 | 9398 |
| 13 | 9979 | 7706 | 14206 | 13455 |
| 14 | 10591 | 8251 | 8689 | 12540 |
| 15 | 10928 | 7284 | 10532 | 12572 |
| 16 | 10268 | 7143 | 8849 | 13842 |
| 17 | 8545 | 9681 | 9978 | 15184 |
| 18 | 8675 | 6859 | 8558 | 11974 |
| 19 | 9854 | 7882 | 8501 | 18233 |
| 20 | 9314 | 8199 | 10532 | 11592 |
| 21 | 9343 | 8872 | 9728 | 12791 |
| 22 | 8105 | 6418 | 10214 | 13301 |
| 23 | 8870 | 7418 | 11012 | 14239 |
| 24 | 9527 | 8764 | 10010 | 12692 |
| 25 | 11072 | 9711 | 13471 | 11316 |

#### 4.1 Results for Baseline Methods Applied to ChIA-Drop Datasets

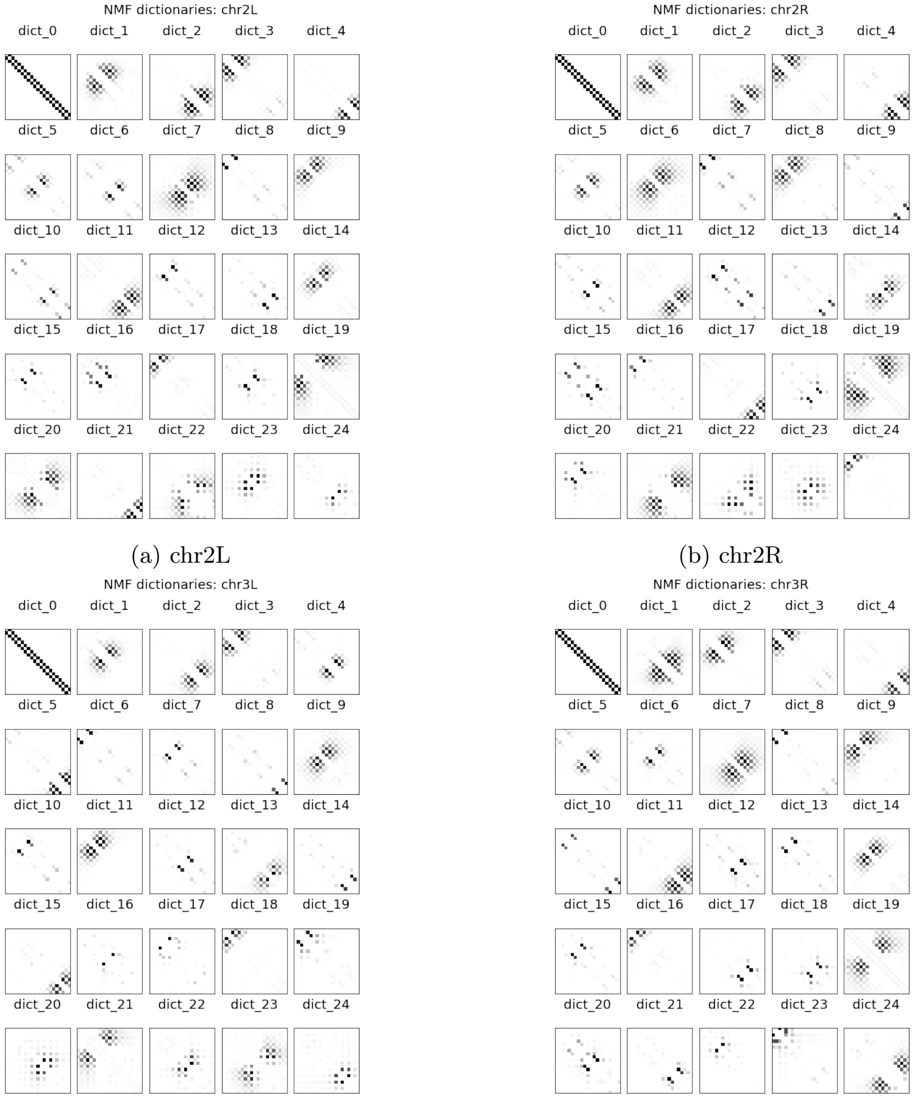


(a) chr2L (b) chr2R

(c) chr3L (d) chr3R

Figure 3: Dictionaries learned by NMF for chr2L, 2R, 3L and 3R.

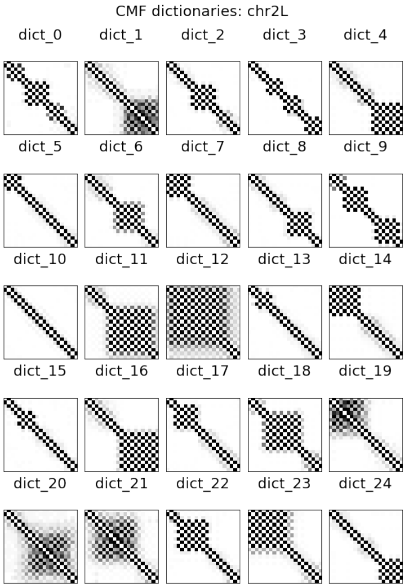


(a) chr2L

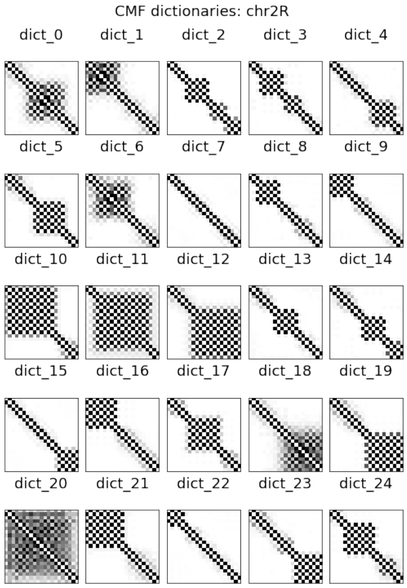


(b) chr2R

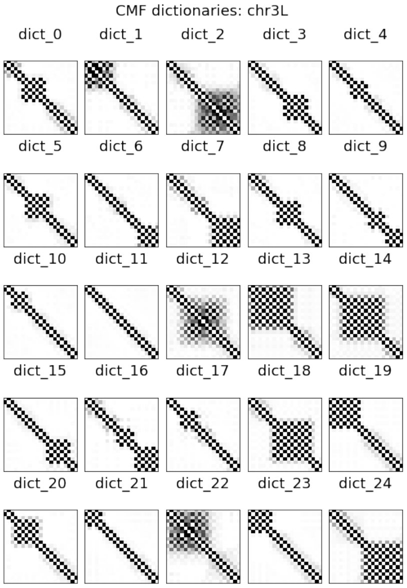


(c) chr3L

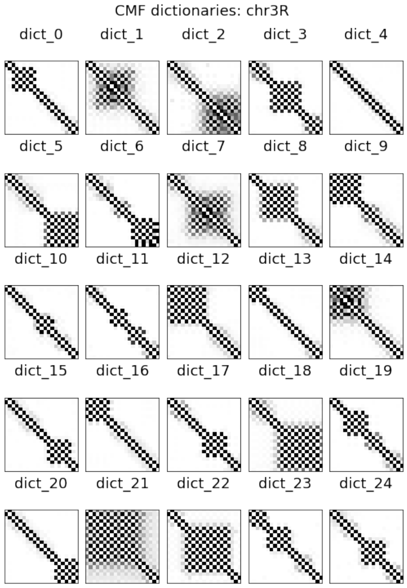


(d) chr3R

Figure 4: Dictionaries learned by CMF for chr2L, 2R, 3L and 3R.

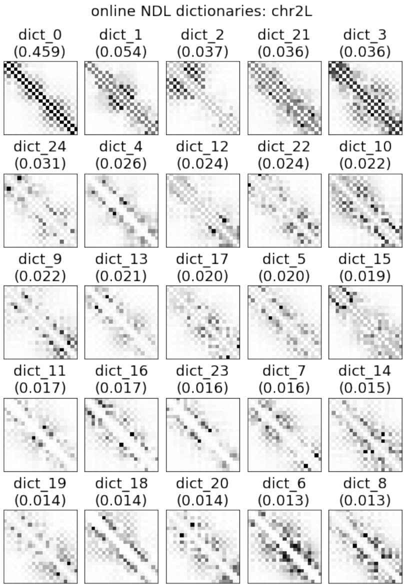


(a) chr2L

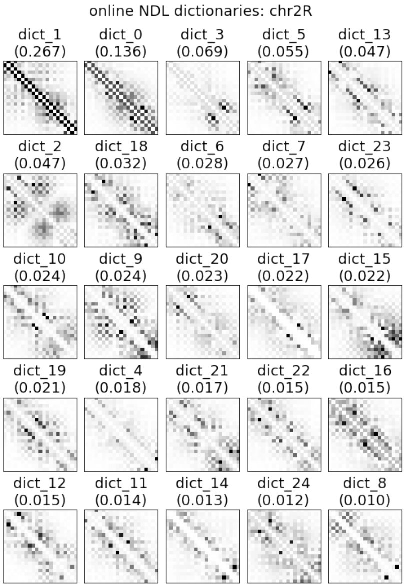


(b) chr2R

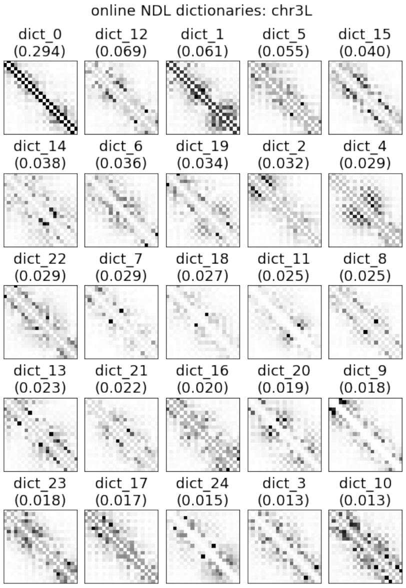


(c) chr3L

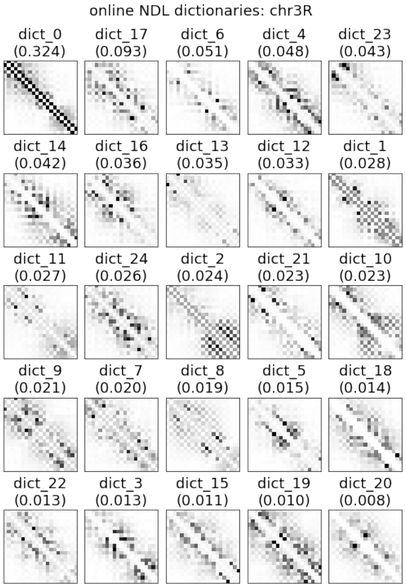


(d) chr3R

Figure 5: Dictionaries learned by online NDL for chr2L, 2R, 3L and 3R.

### 5 Reconstruction of ChIA-Drop Contact Maps

The reconstructions for 4 randomly selected subnetwork samples are shown in Figure 6, providing a means to visually assess the accuracy of reconstructed small-scale interactions.

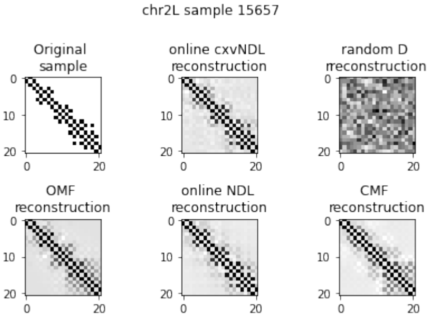


(a) Reconstruction of sample #15657

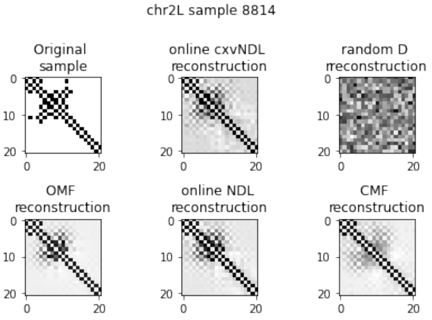


(b) Reconstruction of sample #8814

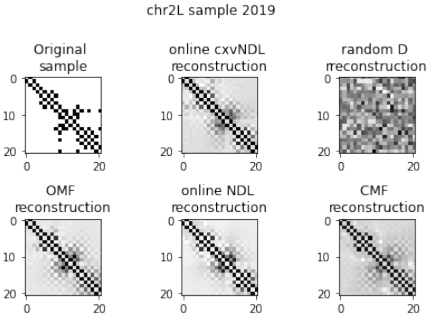


(c) Reconstruction of sample #2019

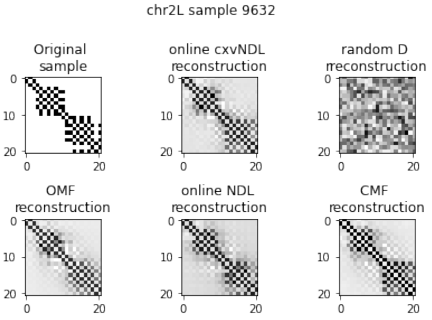


(d) Reconstruction of sample #9632

Figure 6: Reconstructed adjacency matrices for chr2L obtained using different methods and random dictionaries. OMF stands for Ordinary (Standard) MF or NMF.

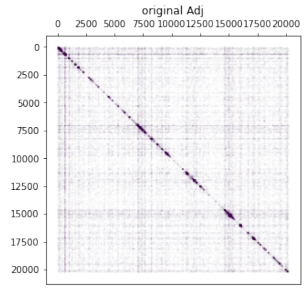


(a) Original adjacency matrix

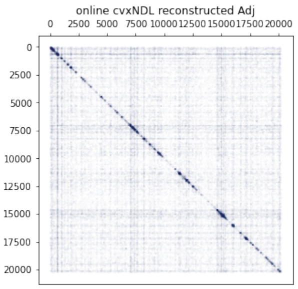


(b) Online cvxNDL

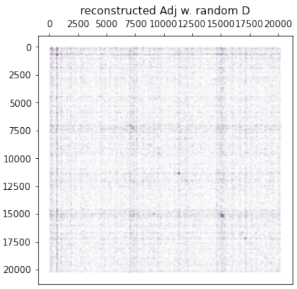


(c) Random dictionaries

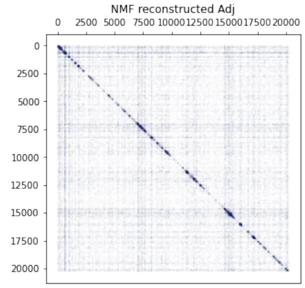


(d) NMF

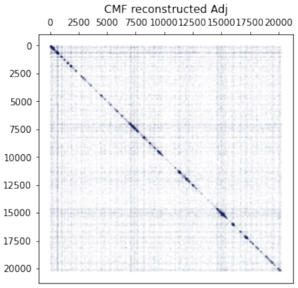


(e) CMF

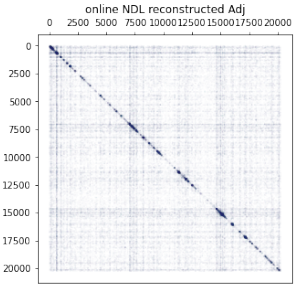


(f) Online NDL

Figure 7: Comparison of network reconstructions obtained using different baseline methods and random dictionaries for *Drosophila* chromosome 2L. (a): The original adjacency matrix; (b, c, d, e, f): Reconstructed network adjacency matrices with online cxvNDL, random dictionary elements, NMF, CMF and online NDL, respectively.

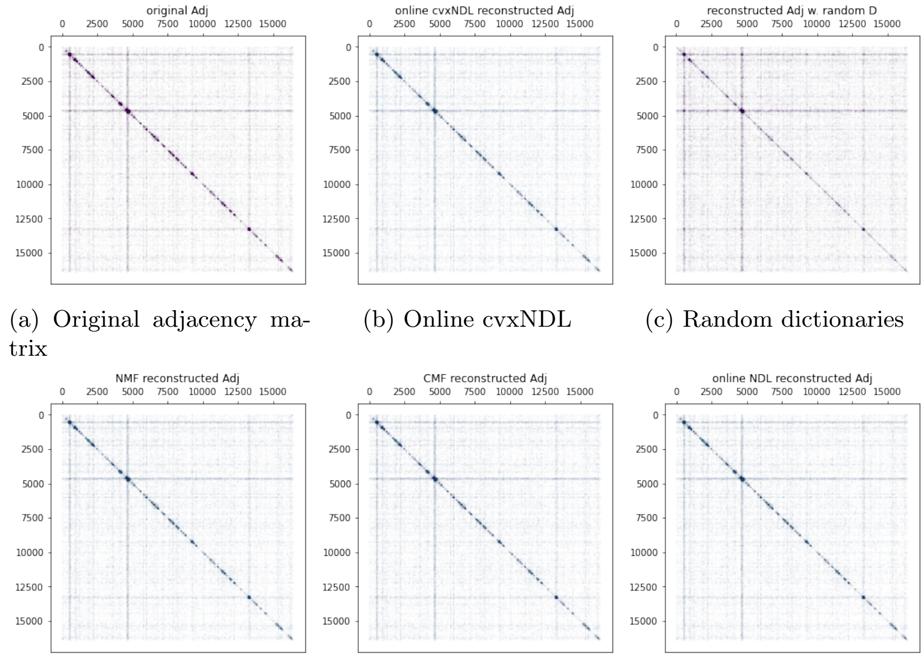


(a) Original adjacency matrix (b) Online cvxNDL (c) Random dictionaries

(d) NMF (e) CMF (f) Online NDL

Figure 8: Reconstructed network comparisons based on different baseline methods and random dictionaries, applied on *Drosophila* chromosome 2R. (a): The original adjacency matrix. (b, c, d, e, f): Reconstructed network adjacency matrices with online cxvNDL, random dictionary elements, NMF, CMF and online NDL.

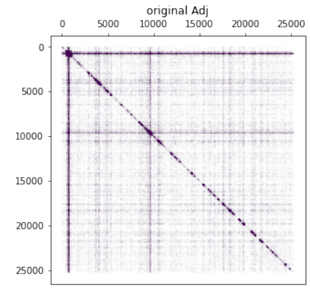


(a) Original adjacency matrix

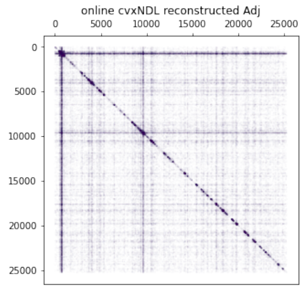


(b) Online cvxNDL

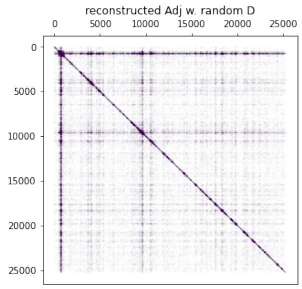


(c) Random dictionaries

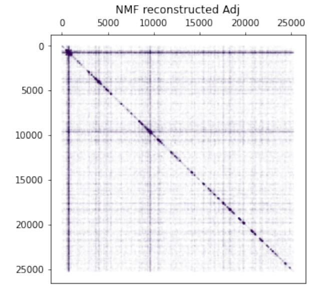


(d) NMF

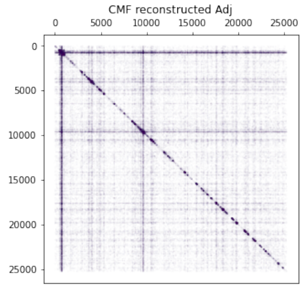


(e) CMF

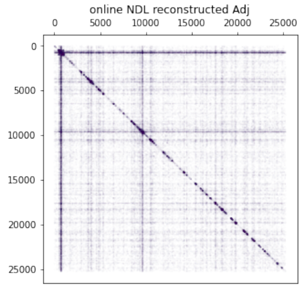


(f) Online NDL

Figure 9: Reconstructed network comparisons based on different baseline methods and random dictionaries, applied on *Drosophila* chromosome 3L. (a): The original adjacency matrix. (b, c, d, e, f): Reconstructed network adjacency matrices with online cxvNDL, random dictionary elements, NMF, CMF and online NDL.

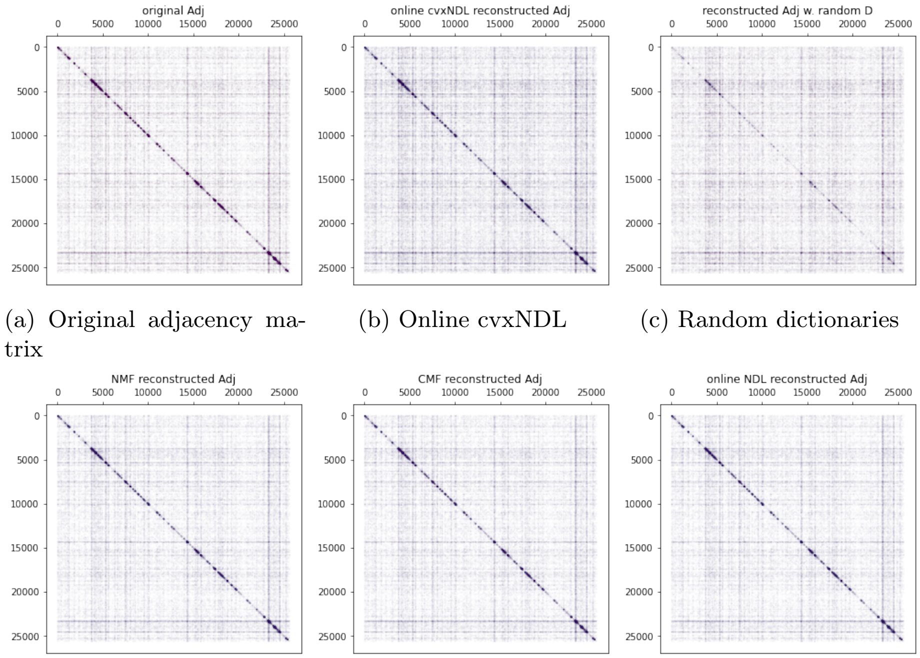


(a) Original adjacency matrix

(b) Online cvxNDL

(c) Random dictionaries

(d) NMF

(e) CMF

(f) Online NDL

Figure 10: Reconstructed network comparisons based on different baseline methods and random dictionaries, applied on *Drosophila* chromosome 3R. (a): The original adjacency matrix. (b, c, d, e, f): Reconstructed network adjacency with online cxvNDL, random dictionary elements, NMF, CMF and online NDL.

### 6 Gene Ontology Enrichment Analysis

To associate a biological function with each dictionary element, we performed a gene ontology (GO) enrichment analysis for each element and the corresponding chromosome. Recall that as a results of the convexity constraint, every dictionary element has its corresponding set of representatives that capture real observed subgraphs which can be mapped back to actual genomic locations. Of most interest is the set of genes that covers at least one vertex in at least one of the representatives, as described in Figure 11.

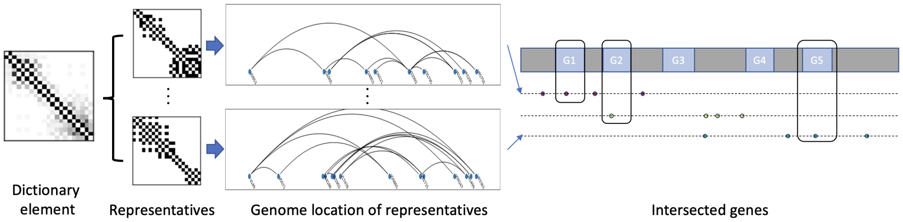


Figure 11: GO enrichment analysis workflow. Each dictionary element is associated with a collection of real subnetwork representatives. These comprise nodes that can be mapped to the genome to identify their locations. A gene is said to cover the node if the genomic fragment corresponding to the node is fully contained within the gene.

Using the set of representative genes, we run the GO enrichment analysis using the annotation category "Biological Process" from `http://geneontology.org`, with the reference list *Drosophila Melanogaster* for each dictionary element. For further analysis, we only selected results with false discovery rate (FDR) $< 0.05$ and hence obtained candidate sets of enriched GO terms. Note that there may be inherently enriched GO terms for each dictionary element due to the sampling bias. To remove this bias, we ran another GO enrichment analysis with all genes on each chromosome and used those results to filter out the background GO terms for each dictionary element.

Furthermore, we utilized the hierarchical structure of GO terms [4], where terms are represented as nodes in a directed acyclic graph, and their relationships are described via arcs in the digraph. A child GO term is considered more specific than a parent GO term. Since the GO graph is not a strict hierarchy (a child node may have multiple parent nodes), to further improve the results, we performed the following processing. For each GO term: i) we first find all the paths between the term and the root node (which is "Biological process" in our setting), and ii) we remove all intermediate parent GO terms from its enriched GO terms set. By iteratively repeating this filtering process for each dictionary element, we derived a set of the most specific GO terms for each dictionary element.

#### 6.1 Dictionary Elements Associated with GO Terms

We investigated the most frequently enriched GO terms as well as the least frequently enriched GO terms for each chromosome and identified the corresponding dictionary elements where they were found to be enriched. The results are shown in Tables 4 to 7. For each dictionary element, we computed its density (complexity) $\rho$ via $\rho = \frac{1}{k^2}\sum_{i,j}\mathbf{D}_{i,j}$ and the median genomic distance between all consecutive pairs of nodes, denoted by $d_{\text{med}}$. The full set of results for the densities and median distances for all dictionary elements and all chromosomes is provided in Tables 2 and 3.

Note that the *Drosophila* S2 cells are embryonic cells, and most GO terms found are related to cellular reproductive process or developmental process, as expected. From the tables, one can also see that different dictionary elements reflect different biological processes and for the same GO term, the dictionary elements share similar patterns. For example, in Table 4, we can see that dictionary elements 19 and 12 share very similar structural patterns, and both of them are enriched in biosynthetic processes of antibacterial peptides. On the other hand, dictionary elements 13 and 8 have a pattern that differs from that of 19 and 12, and they are enriched in dorsal/ventral lineage restriction processes. We also found that dictionary elements with GO term *peripheral nervous system development*, *celluar response to organic substance*, and *neuroblast fate determination* have relatively lower density and smaller median node distances than the top 2 enriched GO terms, *regulation of reproductive process* and *muscle cell cellular homeostasis*. The difference in density and median distance is also reflected by the significantly different dictionary patterns observed, such as dictionary element 12 and dictionary element 5; the former element has a much higher density and median distance than the latter.

There are also a few shared GO terms that are enriched in both chr2L and chr2R (11 shared terms in total) and in both chr3L and chr3R (3 shared terms in total). The results are reported in Table 8 and 9. We found that there are very few shared terms between the two chromosomes when compared to the roughly one hundred uniquely enriched GO terms for each chromosome. Most of the shared terms also have "similar" patterns (which can be seen visually or through a simple computation of the $\ell_2$ distance between their flattened adjacency matrices) of their corresponding dictionary elements.

Table 4: The 5 most and least enriched GO terms within the span of dictionary elements for chr2L. Column '#' indicates the number of dictionary elements that show enrichment for the given GO term. Also reported are up to 3 dictionary elements with the largest importance score in the dictionary, along with the "density" $\rho$ of interactions in the dictionary element and median distance $d_{\text{med}}$ of all adjacent pairs of nodes in its representatives.

| most frequent GO term | # | top 3 dictionaries | least frequent GO term | # | dictionary |
|---|---|---|---|---|---|
| (GO:2000241) regulation of reproductive process | 5 | dict_2 (0.085) 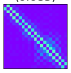 dict_21 (0.070) 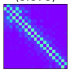 dict_6 (0.044) 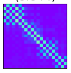 $\rho$=0.134,0.142,0.161 $d_{\text{med}}$=9906,8105,10024 | (GO:0007485) imaginal disc-derived male genitalia development | 1 | dict_21 (0.070) 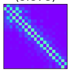 $\rho$=0.142 $d_{\text{med}}$= 8105 |
| (GO:0046716) muscle cell cellular homeostasis | 4 | dict_14 (0.055) 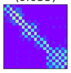 dict_6 (0.044) 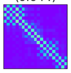 dict_12 (0.029) 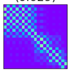 $\rho$=0.141,0.161,0.203 $d_{\text{med}}$=10928,10024,9979 | (GO:0008347) glial cell migration | 1 | dict_5 (0.074) 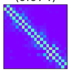 $\rho$=0.132 $d_{\text{med}}$= 8547 |
| (GO:0007422) peripheral nervous system development | 3 | dict_5 (0.074) 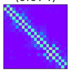 dict_7 (0.061) 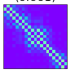 dict_8 (0.057) 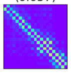 $\rho$=0.132,0.158,0.147 $d_{\text{med}}$=8547,8870,10692 | (GO:0002920) regulation of humoral immune response | 1 | dict_21 (0.070) 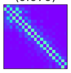 $\rho$=0.142 $d_{\text{med}}$= 8105 |
| (GO:0071310) cellular response to organic substance | 3 | dict_2 (0.085) 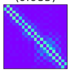 dict_21 (0.070) 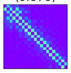 dict_7 (0.061) 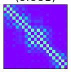 $\rho$=0.134,0.142,0.158 $d_{\text{med}}$=9906,8105,8870 | (GO:0016075) rRNA catabolic process | 1 | dict_8 (0.057) 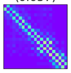 $\rho$=0.147 $d_{\text{med}}$= 10692 |
| (GO:0007400) neuroblast fate determination | 3 | dict_5 (0.074) 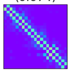 dict_21 (0.070) 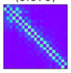 dict_8 (0.057) 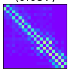 $\rho$=0.132,0.142,0.147 $d_{\text{med}}$=8547,8105,10692 | (GO:0008258) head involution | 1 | dict_8 (0.057) 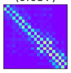 $\rho$=0.147 $d_{\text{med}}$= 10692 |

Table 5: The 5 most and least enriched GO terms within the span of dictionary elements for chr2R. Column '#' indicates the number of dictionary elements that show enrichment for the given GO term. Also reported are up to 3 dictionary elements with the largest importance score in the dictionary, along with the "density" $\rho$ of interactions in the dictionary element and median distance $d_{\text{med}}$ of all adjacent pairs of nodes in its representatives.

| most frequent GO term | # | top 3 dictionaries | least frequent GO term | # | dictionary |
|---|---|---|---|---|---|
| (GO:0030706) germarium-derived oocyte differentiation | 6 | 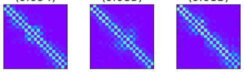  $\rho$=0.140,0.145,0.146 $d_{\text{med}}$=8764,7651,7158 | (GO:0050803) regulation of synapse structure or activity | 1 | 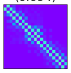  $\rho$=0.140 $d_{\text{med}}$= 8764 |
| (GO:0001700) embryonic development via the syncytial blastoderm | 5 | 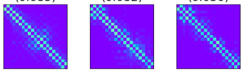  $\rho$=0.145,0.141,0.136 $d_{\text{med}}$=7651,8251,7085 | (GO:0007498) mesoderm development | 1 | 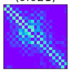  $\rho$=0.183 $d_{\text{med}}$= 7143 |
| (GO:0007451) dorsal/ventral lineage restriction, imaginal disc | 4 | 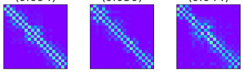  $\rho$=0.140,0.136,0.157 $d_{\text{med}}$=8764,7085,6738 | (GO:0010638) positive regulation of organelle organization | 1 | 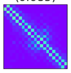  $\rho$=0.145 $d_{\text{med}}$= 7651 |
| (GO:0006964) positive regulation of biosynthetic process of antibacterial peptides active against Gram-negative bacteria | 3 | 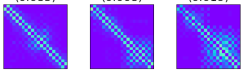  $\rho$=0.145,0.156,0.202 $d_{\text{med}}$=7651,8199,7706 | (GO:0043277) apoptotic cell clearance | 1 | 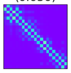  $\rho$=0.136 $d_{\text{med}}$= 7085 |
| (GO:0045476) nurse cell apoptotic process | 3 | 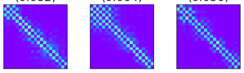  $\rho$=0.141,0.159,0.136 $d_{\text{med}}$=8251,7882,7085 | (GO:0001707) mesoderm formation | 1 | 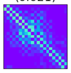  $\rho$=0.183 $d_{\text{med}}$= 7143 |

Table 6: The 5 most and least enriched GO terms within the span of dictionary elements for chr3L. Column '#' indicates the number of dictionary elements that show enrichment for the given GO term. Also reported are up to 3 dictionary elements with the largest importance score in the dictionary, along with the "density" $\rho$ of interactions in the dictionary element and median distance $d_{\text{med}}$ of all adjacent pairs of nodes in its representatives.

| most frequent GO term | # | top 3 dictionaries | least frequent GO term | # | dictionary |
|---|---|---|---|---|---|
| (GO:0009631) cold acclimation | 2 | 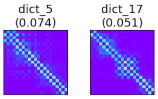<br>$\rho$=0.148,0.152<br>$d_{\text{med}}$=10608,8558 | (GO:0035070) salivary gland histolysis | 1 | 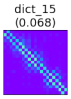<br>$\rho$=0.143<br>$d_{\text{med}}$= 8849 |
| (GO:0009408) response to heat | 2 | 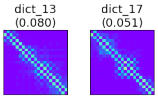<br>$\rho$=0.147,0.152<br>$d_{\text{med}}$=8689,8558 | (GO:0046843) dorsal appendage formation | 1 | 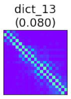<br>$\rho$=0.147<br>$d_{\text{med}}$= 8689 |
| (GO:0007616) long-term memory | 2 | 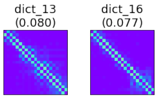<br>$\rho$=0.147,0.126<br>$d_{\text{med}}$=8689,9978 | (GO:0007097) nuclear migration | 1 | 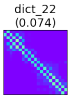<br>$\rho$=0.134<br>$d_{\text{med}}$= 11012 |
| (GO:0061077) chaperone-mediated protein folding | 2 | 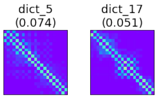<br>$\rho$=0.148,0.152<br>$d_{\text{med}}$=10608,8558 | (GO:0035071) salivary gland cell autophagic cell death | 1 | 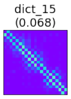<br>$\rho$=0.143<br>$d_{\text{med}}$= 8849 |
| (GO:0008587) imaginal disc-derived wing margin morphogenesis | 2 | 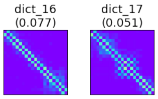<br>$\rho$=0.126,0.152<br>$d_{\text{med}}$=9978,8558 | (GO:0007528) neuromuscular junction development | 1 | 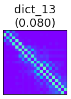<br>$\rho$=0.147<br>$d_{\text{med}}$= 8689 |

Table 7: The 5 most and least enriched GO terms within the span of dictionary elements for chr3R. Column '#' indicates the number of dictionary elements that show enrichment for the given GO term. Also reported are up to 3 dictionary elements with the largest importance score in the dictionary, along with the "density" $\rho$ of interactions in the dictionary element and median distance $d_{\mathrm{med}}$ of all adjacent pairs of nodes in its representatives.

| most frequent GO term | # | top 3 dictionaries | least frequent GO term | # | dictionary |
|---|---|---|---|---|---|
| (GO:0001819) positive regulation of cytokine production | 7 | 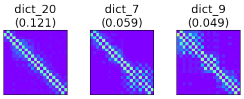  $\rho$=0.126,0.146,0.157 $d_{\mathrm{med}}$=12791,12830,11930 | (GO:0061448) connective tissue development | 1 | 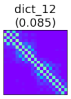  $\rho$=0.142 $d_{\mathrm{med}}$= 13455 |
| (GO:0008015) blood circulation | 7 | 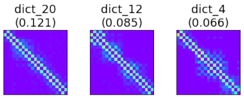  $\rho$=0.126,0.142,0.138 $d_{\mathrm{med}}$=12791,13455,13674 | (GO:0051282) regulation of sequestering of calcium ion | 1 | 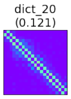  $\rho$=0.126 $d_{\mathrm{med}}$= 12791 |
| (GO:0045948) positive regulation of translational initiation | 5 | 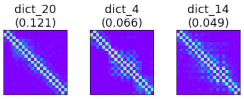  $\rho$=0.126,0.138,0.162 $d_{\mathrm{med}}$=12791,13674,12572 | (GO:0043123) positive regulation of I-kappaB kinase/NF-kappaB signaling | 1 | 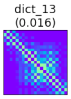  $\rho$=0.204 $d_{\mathrm{med}}$= 12540 |
| (GO:0042177) negative regulation of protein catabolic process | 5 | 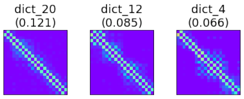  $\rho$=0.126,0.142,0.138 $d_{\mathrm{med}}$=12791,13455,13674 | (GO:0007435) salivary gland morphogenesis | 1 | 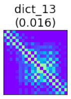  $\rho$=0.204 $d_{\mathrm{med}}$= 12540 |
| (GO:0043065) positive regulation of apoptotic process | 4 | 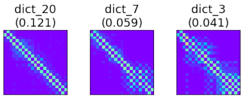  $\rho$=0.126,0.146,0.179 $d_{\mathrm{med}}$=12791,12830,11748 | (GO:0045738) negative regulation of DNA repair | 1 | 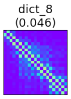  $\rho$=0.183 $d_{\mathrm{med}}$= 12493 |

Table 8: GO terms shared between chr2L and chr2R.

| GO_term | chr2L dictionaries | chr2R dictionaries |
|---|---|---|
| (GO:0016325) oocyte microtubule cytoskeleton organization | dict_5 (0.074), dict_7 (0.061), dict_6 (0.044) 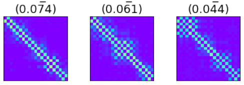 | dict_14 (0.013) 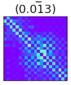 |
| (GO:1901701) cellular response to oxygen-containing compound | dict_2 (0.085), dict_7 (0.061) 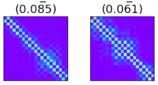 | dict_8 (0.050) 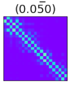 |
| (GO:0007298) border follicle cell migration | dict_2 (0.085), dict_21 (0.070) 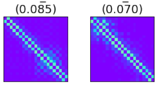 | dict_4 (0.085), dict_3 (0.083), dict_18 (0.064) 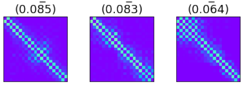 |
| (GO:0043410) positive regulation of MAPK cascade | dict_2 (0.085), dict_8 (0.057) 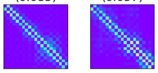 | dict_4 (0.085), dict_8 (0.050) 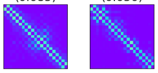 |
| (GO:0016049) cell growth | dict_21 (0.070) 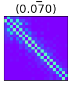 | dict_8 (0.050) 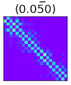 |
| (GO:0035331) negative regulation of hippo signaling | dict_8 (0.057) 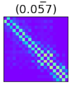 | dict_4 (0.085) 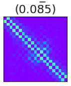 |
| (GO:0051962) positive regulation of nervous system development | dict_7 (0.061) 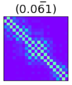 | dict_15 (0.021) 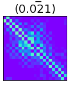 |
| (GO:0060322) head development | dict_8 (0.057) 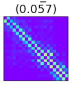 | dict_4 (0.085) 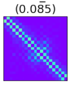 |
| (GO:0007293) germarium-derived egg chamber formation | dict_8 (0.057) 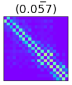 | dict_23 (0.094), dict_4 (0.085), dict_13 (0.082), dict_15 (0.021) 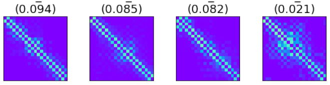 |
| (GO:0002164) larval development | dict_6 (0.044) 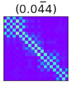 | dict_15 (0.021) 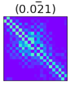 |
| (GO:0007420) brain development | dict_6 (0.044) 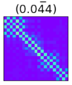 | dict_4 (0.085), dict_18 (0.064) 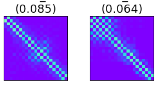 |

Table 9: GO terms shared between chr3L and chr3R.

| GO_term | chr3L dictionaries | chr3R dictionaries |
|---|---|---|
| (GO:0070373) negative regulation of ERK1 and ERK2 cascade | dict_13 (0.080) 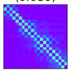 dict_22 (0.074) 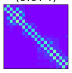 dict_3 (0.045) 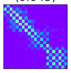 dict_1 (0.035) 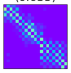 | dict_8 (0.046) 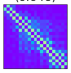 |
| (GO:0007140) male meiotic nuclear division | dict_23 (0.029) 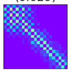 | dict_24 (0.017) 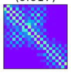 |
| (GO:0046777) protein autophosphorylation | dict_22 (0.074) 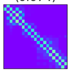 | dict_8 (0.046) 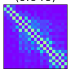 |

#### 6.2 Additional Results

Here we report more detailed results for each dictionary element, including its number of enriched GO terms and importance scores (Tables 10, 11, 12, 13).

Table 10: Number of enriched GO terms for each dictionary element identified for chr2L.

| | # GO terms | | # GO terms | | # GO terms | | # GO terms | | # GO terms |
|---|---|---|---|---|---|---|---|---|---|
| dict_0 (0.077) 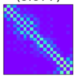 | 2 | dict_5 (0.074) 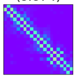 | 15 | dict_10 (0.018) 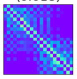 | 0 | dict_15 (0.038) 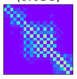 | 0 | dict_20 (0.024) 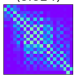 | 0 |
| dict_1 (0.019) 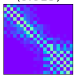 | 0 | dict_6 (0.044) 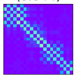 | 19 | dict_11 (0.022) 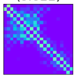 | 2 | dict_16 (0.030) 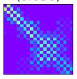 | 2 | dict_21 (0.070) 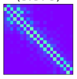 | 27 |
| dict_2 (0.085) 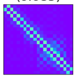 | 20 | dict_7 (0.061) 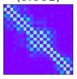 | 24 | dict_12 (0.029) 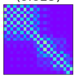 | 1 | dict_17 (0.045) 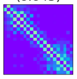 | 0 | dict_22 (0.046) 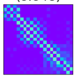 | 1 |
| dict_3 (0.030) 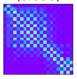 | 0 | dict_8 (0.057) 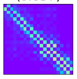 | 31 | dict_13 (0.014) 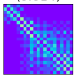 | 0 | dict_18 (0.030) 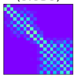 | 0 | dict_23 (0.014) 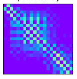 | 0 |
| dict_4 (0.059) 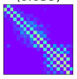 | 0 | dict_9 (0.017) 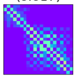 | 0 | dict_14 (0.055) 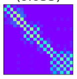 | 6 | dict_19 (0.016) 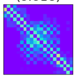 | 0 | dict_24 (0.025) 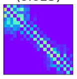 | 0 |

Table 11: Number of enriched GO terms for each dictionary element identified for chr2R.

| # GO terms | # GO terms | # GO terms | # GO terms | # GO terms |
|---|---|---|---|---|
| dict_0 (0.044) 4 | dict_5 (0.014) 0 | dict_10 (0.014) 0 | dict_15 (0.021) 23 | dict_20 (0.041) 6 |
| dict_1 (0.041) 1 | dict_6 (0.025) 0 | dict_11 (0.042) 1 | dict_16 (0.018) 0 | dict_21 (0.019) 0 |
| dict_2 (0.035) 0 | dict_7 (0.037) 1 | dict_12 (0.019) 2 | dict_17 (0.020) 0 | dict_22 (0.019) 8 |
| dict_3 (0.083) 12 | dict_8 (0.050) 17 | dict_13 (0.082) 9 | dict_18 (0.064) 8 | dict_23 (0.094) 10 |
| dict_4 (0.085) 40 | dict_9 (0.030) 0 | dict_14 (0.013) 5 | dict_19 (0.068) 7 | dict_24 (0.022) 2 |

Table 12: Number of enriched GO terms for each dictionary element identified for chr3L.

| # GO terms | # GO terms | # GO terms | # GO terms | # GO terms |
|---|---|---|---|---|
| dict_0 (0.022) 0 | dict_5 (0.074) 6 | dict_10 (0.023) 2 | dict_15 (0.068) 10 | dict_20 (0.025) 0 |
| dict_1 (0.035) 3 | dict_6 (0.028) 1 | dict_11 (0.027) 0 | dict_16 (0.077) 14 | dict_21 (0.018) 0 |
| dict_2 (0.049) 0 | dict_7 (0.029) 1 | dict_12 (0.021) 1 | dict_17 (0.051) 9 | dict_22 (0.074) 4 |
| dict_3 (0.045) 3 | dict_8 (0.020) 0 | dict_13 (0.080) 16 | dict_18 (0.023) 4 | dict_23 (0.029) 3 |
| dict_4 (0.074) 3 | dict_9 (0.023) 0 | dict_14 (0.009) 0 | dict_19 (0.037) 0 | dict_24 (0.040) 0 |

Table 13: Number of enriched GO terms for each dictionary element identified for chr3R.

| | # GO terms | | # GO terms | | # GO terms | | # GO terms | | # GO terms |
|---|---|---|---|---|---|---|---|---|---|
| dict_0 (0.046) 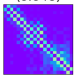 | 15 | dict_5 (0.038) 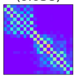 | 2 | dict_10 (0.040) 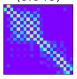 | 5 | dict_15 (0.016) 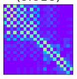 | 8 | dict_20 (0.121) 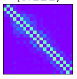 | 124 |
| dict_1 (0.042) 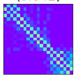 | 9 | dict_6 (0.029) 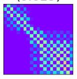 | 2 | dict_11 (0.021) 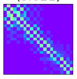 | 0 | dict_16 (0.019) 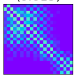 | 0 | dict_21 (0.041) 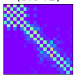 | 10 |
| dict_2 (0.062) 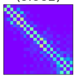 | 13 | dict_7 (0.059) 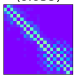 | 14 | dict_12 (0.085) 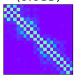 | 16 | dict_17 (0.015) 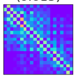 | 0 | dict_22 (0.022) 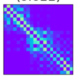 | 4 |
| dict_3 (0.041) 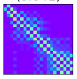 | 7 | dict_8 (0.046) 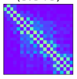 | 25 | dict_13 (0.016) 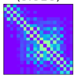 | 57 | dict_18 (0.014) 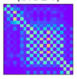 | 0 | dict_23 (0.016) 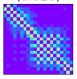 | 0 |
| dict_4 (0.066) 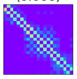 | 20 | dict_9 (0.049) 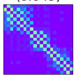 | 1 | dict_14 (0.049) 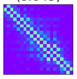 | 6 | dict_19 (0.027) 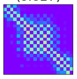 | 0 | dict_24 (0.017) 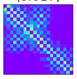 | 4 |

### 7 RNA-Seq Coexpression Analysis

The ChIA-Drop dataset [5] used for learning dictionaries of chromatin interactions lacks RNA-Seq replicates, posing a challenge when trying to validate our results through coexpression analysis. To address this limitation, we retrieved RNA-Seq data corresponding to untreated S2 cell lines of *Drosophila Melanogaster* from the Digital Expression Explorer (DEE2) repository. DEE2 provides uniformly processed RNA-Seq data sourced from the publicly available NCBI Sequence Read Archive (SRA) [6]. In total, we retrieved 20 samples from untreated S2 cell lines with their IDs reported in Table 14.

Table 14: Sample IDs retrieved from NCBI Sequence Read Archive for RNA-Seq coexpression analysis.

| | | | | |
|---|---|---|---|---|
| SRR12191916 | SRR12191917 | SRR12191918 | SRR12191920 | SRR12191921 |
| SRR12191923 | SRR12191927 | SRR2442878 | SRR2442879 | SRR3065067 |
| SRR5340065 | SRR5340066 | SRR5340069 | SRR5340070 | SRR5340071 |
| SRR5340072 | SRR6930637 | SRR8108628 | SRR8108629 | SRR8108630 |

To ensure consistent normalization across all samples, we use the trimmed mean of M values (TMM) method [7], available through the edgeR package [8]. This is of crucial importance when jointly analyzing samples from multiple sources. We selected the most relevant genes by filtering the list of covered genes and retaining only those with more than 95% overlap with the gene promoter regions, as defined in the *Ensmbl* browser. Subsequently, for each dictionary element, we collected all genes covered by it and calculated the pairwise Pearson correlation coefficient of expressions of pairs of genes in the set. For a pair of random variables $X_1$ and $X_2$, the correlation coefficient is defined as

$$\rho_{X_1X_2} = \frac{\text{Covariance}(X_1, X_2)}{\text{Var}(X_1)\text{Var}(X_2)}$$

For two genes $G_1$ and $G_2$, let $X_1$ and $X_2$ be vectors of normalized read counts. The Pearson correlation coefficient can be written as

$$\rho_{G_1G_2} = \frac{\sum_{i=1}^{n}(x_{1i} - \bar{x}_1)(x_{2i} - \bar{x}_2)}{\sqrt{\sum_{i=1}^{n}(x_{1i} - \bar{x}_1)^2}\sqrt{\sum_{i=1}^{n}(x_{2i} - \bar{x}_2)^2}}$$

where

$n$ is the number of samples,

$\bar{x}_1 = \frac{\sum_{i=1}^{n} x_{1i}}{n}$ and $\bar{x}_2 = \frac{\sum_{i=1}^{n} x_{2i}}{n}$ are sample means.

To visualize the underlying coexpression clusters within the genes, we performed hierarchical clustering. We report the mean correlation statistics as well as mean statistics for positively correlated genes for each dictionary element. Correlation plots for all dictionary elements are shown in Figures 12, 13, 14 and 15.

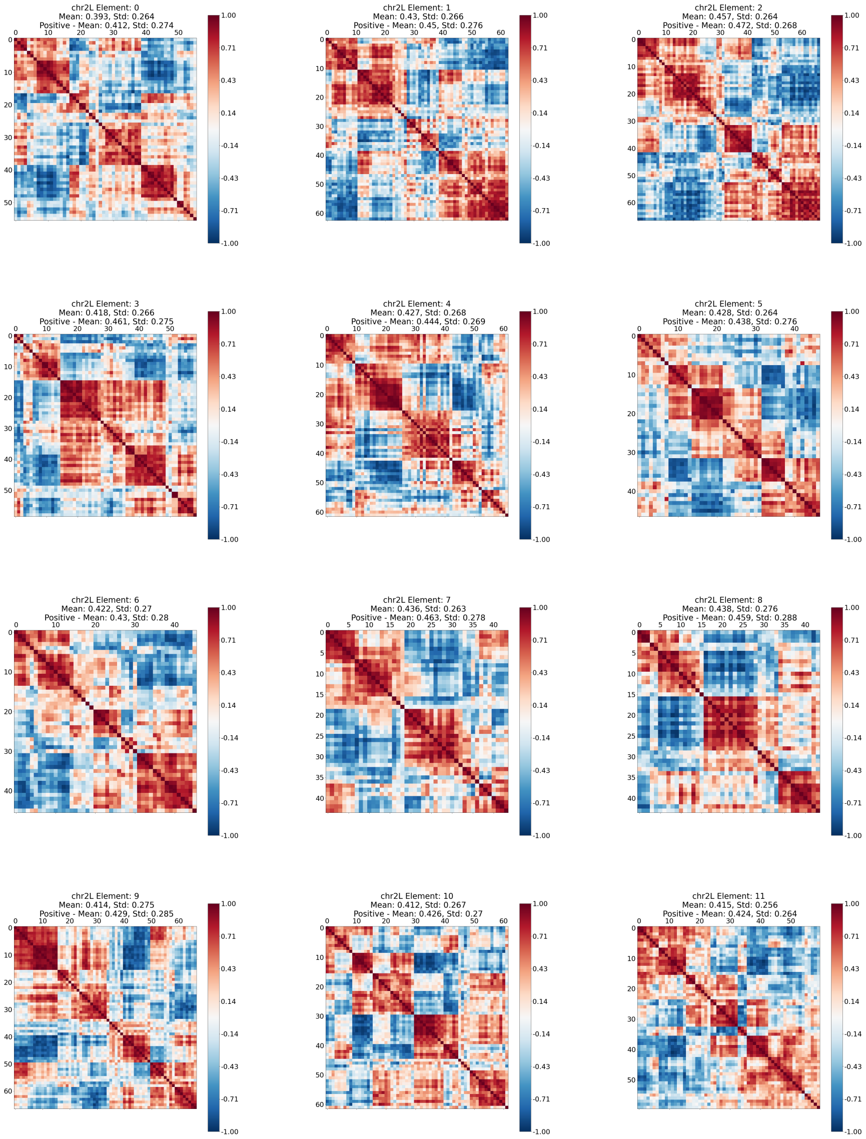


Figure 12: Pairwise coexpression of genes covered by various dictionary elements for chr 2L obtained through online cvxNDL. We calculated the mean and standard deviation of absolute pairwise coexpression values, along with the mean and standard deviation of coexpression values specifically for all positively correlated gene pairs.

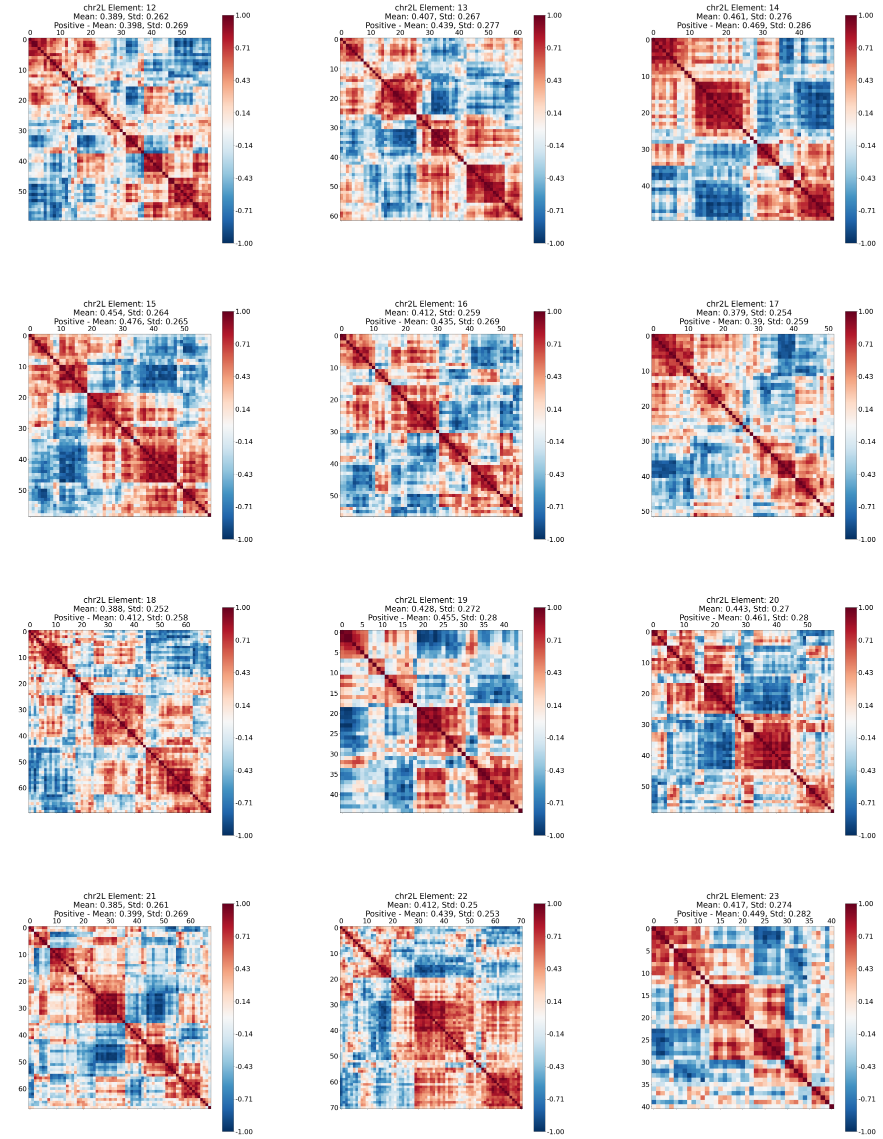


Figure 12: Pairwise coexpression of genes covered by various dictionary elements for chr 2L obtained through online cvxNDL. We calculated the mean and standard deviation of absolute pairwise coexpression values, along with the mean and standard deviation of coexpression values specifically for all positively correlated gene pairs.

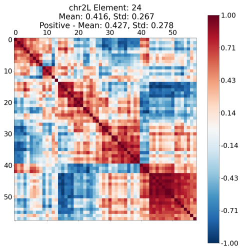


Figure 12: Pairwise coexpression of genes covered by various dictionary elements for chr 2L obtained through online cvxNDL. We calculated the mean and standard deviation of absolute pairwise coexpression values, along with the mean and standard deviation of coexpression values specifically for all positively correlated gene pairs.

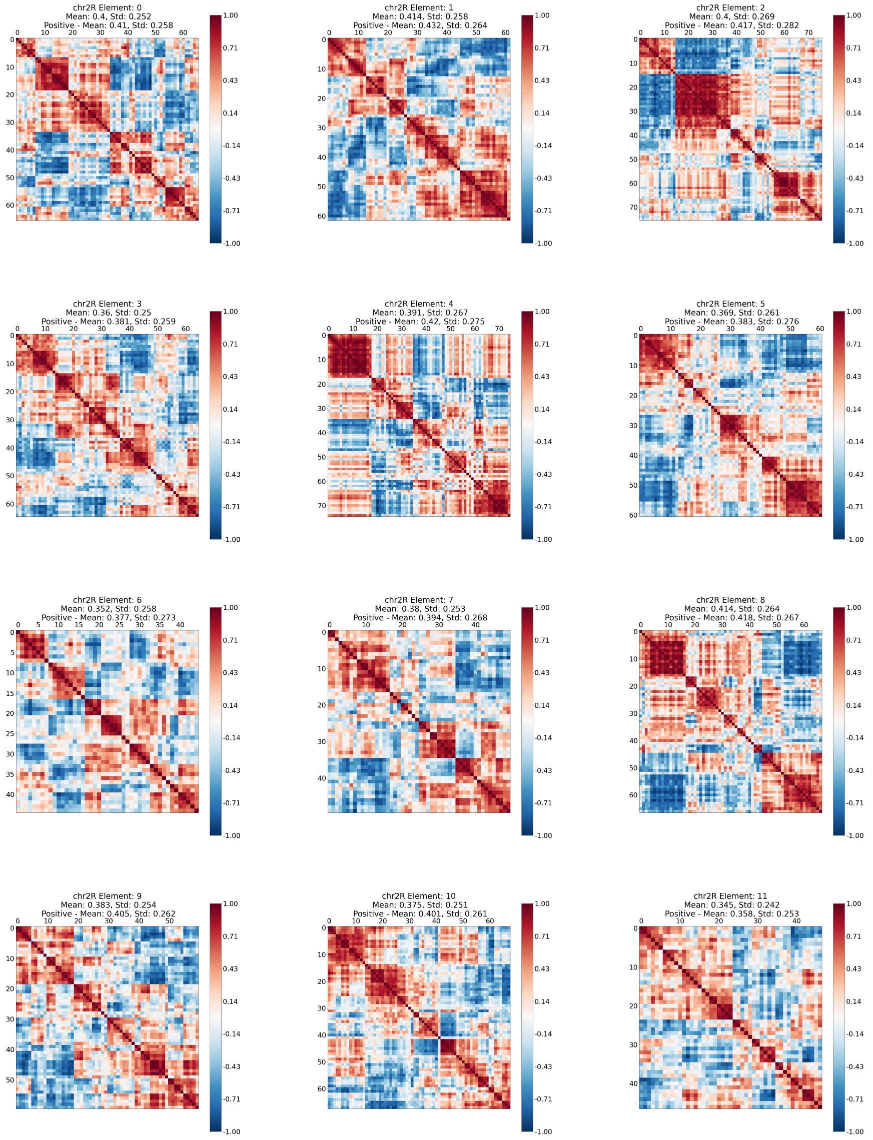


Figure 13: Pairwise coexpression of genes covered by various dictionary elements for chr 2R obtained through online cvxNDL. We calculated the mean and standard deviation of absolute pairwise coexpression values, along with the mean and standard deviation of coexpression values specifically for all positively correlated gene pairs.

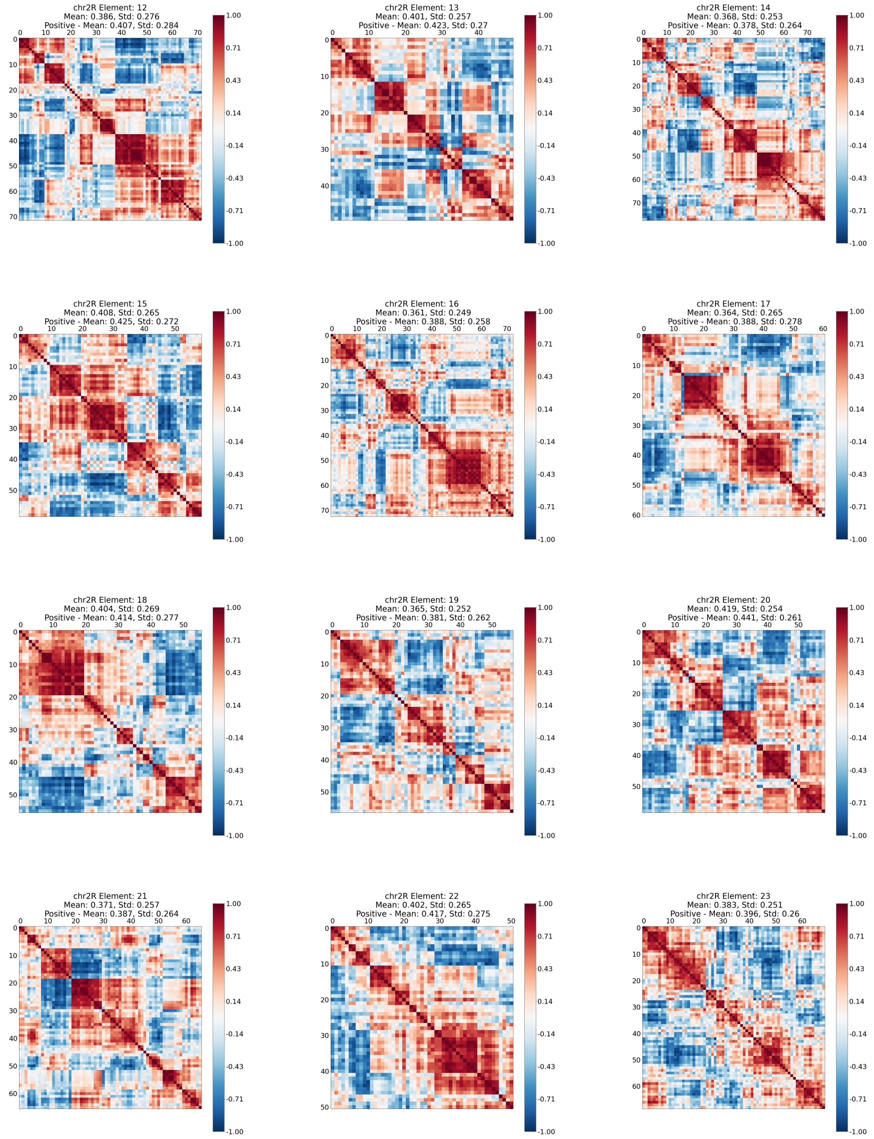


Figure 13: Pairwise coexpression of genes covered by various dictionary elements for chr 2R obtained through online cvxNDL. We calculated the mean and standard deviation of absolute pairwise coexpression values, along with the mean and standard deviation of coexpression values specifically for all positively correlated gene pairs.

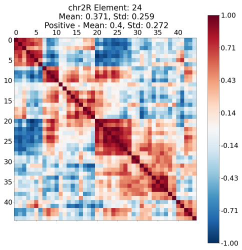


Figure 13: Pairwise coexpression of genes covered by various dictionary elements for chr 2R obtained through online cvxNDL. We calculated the mean and standard deviation of absolute pairwise coexpression values, along with the mean and standard deviation of coexpression values specifically for all positively correlated gene pairs.

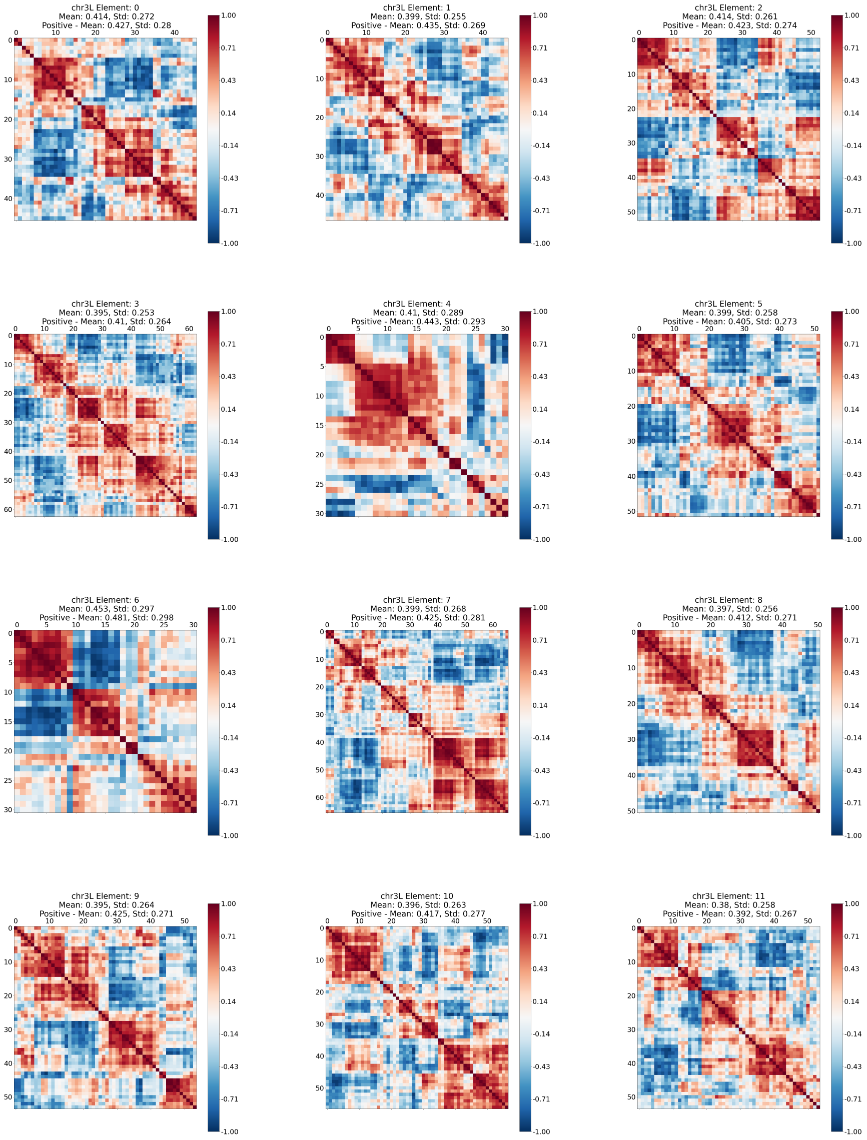


Figure 14: Pairwise coexpression of genes covered by various dictionary elements for chr 3L obtained through online cvxNDL. We calculated the mean and standard deviation of absolute pairwise coexpression values, along with the mean and standard deviation of coexpression values specifically for all positively correlated gene pairs.

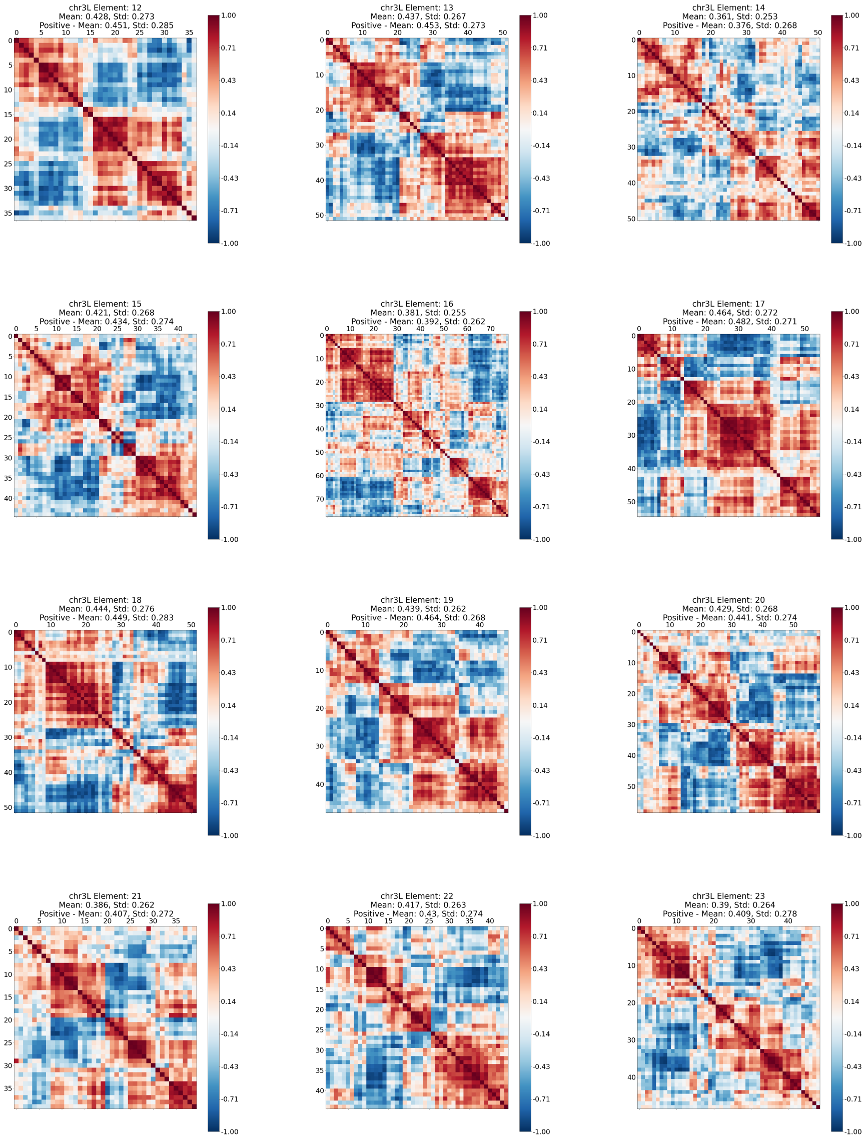


Figure 14: Pairwise coexpression of genes covered by various dictionary elements for chr 3L obtained through online cvxNDL. We calculated the mean and standard deviation of absolute pairwise coexpression values, along with the mean and standard deviation of coexpression values specifically for all positively correlated gene pairs.

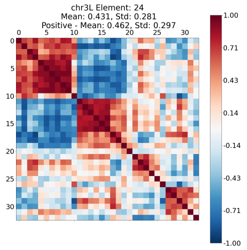


Figure 14: Pairwise coexpression of genes covered by various dictionary elements for chr 3L obtained through online cvxNDL. We calculated the mean and standard deviation of absolute pairwise coexpression values, along with the mean and standard deviation of coexpression values specifically for all positively correlated gene pairs.

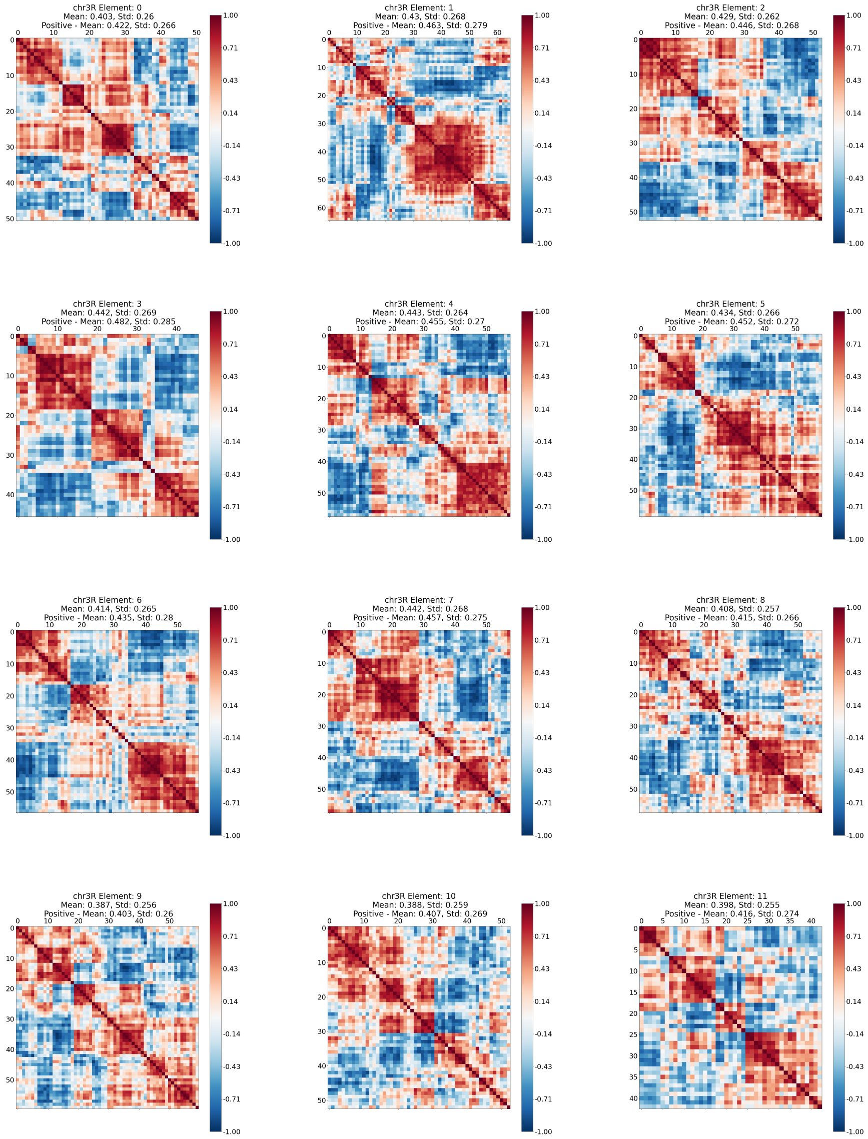


Figure 15: Pairwise coexpression of genes covered by various dictionary elements for chr 3R obtained through online cvxNDL. We calculated the mean and standard deviation of absolute pairwise coexpression values, along with the mean and standard deviation of coexpression values specifically for all positively correlated gene pairs.

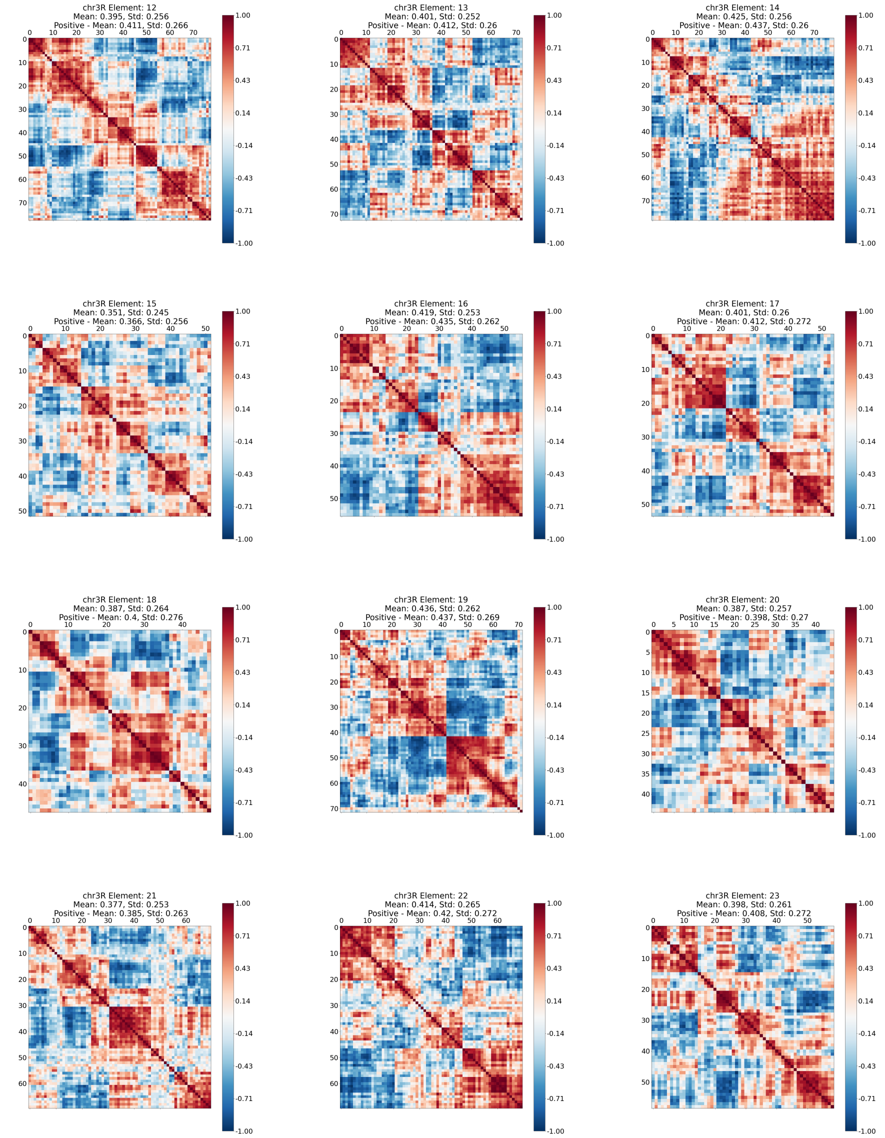


Figure 15: Pairwise coexpression of genes covered by various dictionary elements for chr 3R obtained through online cvxNDL. We calculated the mean and standard deviation of absolute pairwise coexpression values, along with the mean and standard deviation of coexpression values specifically for all positively correlated gene pairs.

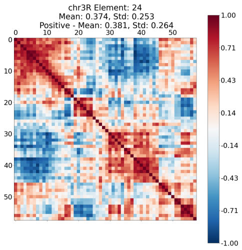


Figure 15: Pairwise coexpression of genes covered by various dictionary elements for chr 3R obtained through online cvxNDL. We calculated the mean and standard deviation of absolute pairwise coexpression values, along with the mean and standard deviation of coexpression values specifically for all positively correlated gene pairs.

### 8 STRING interaction network and FlyMine

The STRING interaction network [9] provides a confidence score indicating the interaction likelihood between a pair of proteins within an organism. This score reflects both direct interactions via physical protein binding and indirect interactions by virtue of the proteins participating in the same cellular pathways. The confidence level of interaction between a pair of proteins can vary from 0, indicating very low confidence, to 1000, indicating very high confidence. Figure 16a shows the distribution of confidence levels between all pairs of proteins in the STRING database for *Drosophila Melanogaster*. A large majority of these interactions are very low confidence. To focus on more reliable interactions, we filtered the protein interactions to retain only those with a confidence score exceeding 200, resulting in a refined dataset shown in Figure 16b. By mapping these proteins back to their corresponding genes, we derived an induced network representing gene-gene interactions.

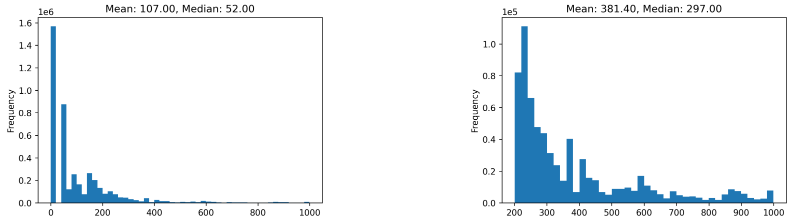


(a) All confidence values.

(b) Confidence values filtered for > 200.

Figure 16: Histogram of confidence values for pairwise interaction of proteins in the STRING interaction network for *Drosophila Melanogaster*.

For the online cvxNDL dictionary, we calculated the mean confidence level for all pairs of proteins. We also repeated the same experiments with a randomly constructed dictionary as a control. Figure 17 shows the mean confidence level and confidence interval for a subset of dictionary elements. We performed a K-S test with the null hypothesis that the two sets of confidence scores for pairwise interactions belonging to online cvxNDL dictionaries and randomly constructed dictionaries are drawn from the same distribution. We rejected the null hypothesis with p-value $< 0.05$.

Flymine [10] is a large genomic and proteomic database for *Drosophila Melanogaster*. We used FlyMine to retrieve a list of upregulated genes in S2 cell lines. We observe that the upregulated genes are overrepresented in our dictionary elements. To test our hypothesis, we performed the hypergeometric overrepresentation test. Our null hypothesis is that the proportion of upregulated genes in our dictionary elements is no higher than the overall proportion of upregulated genes in S2 cell lines. We rejected the null hypothesis (p-value $< 0.05$) for all dictionary elements for all chromosomes except a small subset of 4 dictionary elements (1 dictionary element from chr2R and 3 dictionary elements

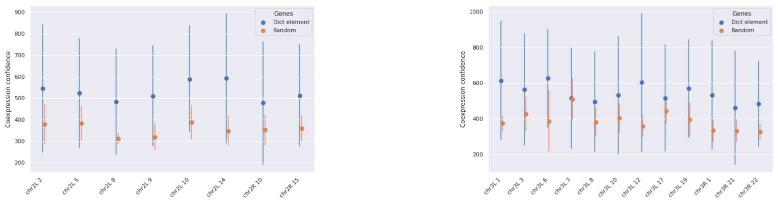


(a) Mean confidence value for dictionary elements from chr2L and chr2R.

(b) Mean confidence value for dictionary elements from chr3L and chr3R

Figure 17: Confidence levels for pairwise interaction of proteins for dictionary elements based on STRING interaction network.

from chr3L). The p-values for all dictionary elements are shown in Table 15.

Table 15: Results for hypergeometric overrepresentation test for all dictionary elements. We report the p-values corresponding to the null hypothesis that the proportion of upregulated genes in our dictionary elements is no higher than the overall proportion of upregulated genes in S2 cell lines.

| dictionary element | chr2L | chr2R | chr3L | chr3R |
|---|---|---|---|---|
| 0 | 1.18E-03 | 5.90E-07 | 7.96E-05 | 3.24E-05 |
| 1 | 1.93E-08 | 8.13E-06 | 5.38E-04 | 9.23E-09 |
| 2 | 4.36E-08 | 4.44E-07 | 1.40E-03 | 1.36E-02 |
| 3 | 8.13E-06 | 7.92E-05 | 1.65E-04 | 4.49E-08 |
| 4 | 4.50E-06 | 1.83E-04 | 2.54E-03 | 4.88E-12 |
| 5 | 1.23E-06 | 3.93E-04 | 3.53E-03 | 5.84E-05 |
| 6 | 1.26E-03 | 2.88E-03 | 5.58E-03 | 6.07E-06 |
| 7 | 1.60E-03 | 3.88E-06 | 1.76E-03 | 1.39E-05 |
| 8 | 3.50E-05 | 9.15E-07 | 1.22E-04 | 3.03E-05 |
| 9 | 2.17E-04 | 2.17E-06 | 2.73E-04 | 4.36E-07 |
| 10 | 1.02E-05 | 3.57E-02 | 5.23E-06 | 2.37E-06 |
| 11 | 1.82E-05 | 8.94E-04 | 8.92E-02 | 1.96E-04 |
| 12 | 2.08E-06 | 8.90E-04 | 2.01E-01 | 3.23E-05 |
| 13 | 8.12E-05 | 8.52E-03 | 3.40E-05 | 1.73E-04 |
| 14 | 1.95E-05 | 1.41E-04 | 1.93E-03 | 1.84E-10 |
| 15 | 6.95E-08 | 5.78E-05 | 1.20E-02 | 8.32E-05 |
| 16 | 5.02E-03 | 7.60E-04 | 1.78E-03 | 4.82E-06 |
| 17 | 3.24E-04 | 5.41E-02 | 9.17E-06 | 7.53E-04 |
| 18 | 1.78E-03 | 6.04E-06 | 1.96E-02 | 3.89E-06 |
| 19 | 3.89E-04 | 3.56E-05 | 8.10E-04 | 6.86E-08 |
| 20 | 1.75E-08 | 2.90E-04 | 5.02E-03 | 1.50E-04 |
| 21 | 6.41E-03 | 1.55E-02 | 3.72E-06 | 8.88E-10 |
| 22 | 2.99E-03 | 1.40E-03 | 2.24E-05 | 9.23E-09 |
| 23 | 1.65E-05 | 6.78E-03 | 5.98E-03 | 3.42E-07 |
| 24 | 2.54E-06 | 1.03E-04 | 6.22E-02 | 7.19E-08 |
